# A novel allele of *Mitogen Activated Protein Kinase 6* is linked to disease resistance and constitutive immunity in rice

**DOI:** 10.1101/2025.03.25.645290

**Authors:** Gokulan C.G., Mohammed Jamaloddin, Deepti Rao, Namami Gaur, Amita Rani Supriyo, Nisha Sao, Deepak Niranjan, Shrish Tiwari, Gouri S. Laha, Kalyani M. Barbadikar, Raman Meenakshi Sundaram, Sheshu Madhav Maganti, Hitendra K. Patel, Ramesh V. Sonti

## Abstract

Bacterial Blight (BB) is the most serious bacterial disease of rice. Usage of resistant cultivars has been a successful strategy in controlling BB. However, constant selection pressure causes the evolution of hypervirulent pathogenic strains that break down resistance, thus necessitating additional sources of resistance. An induced-mutant line derived from the elite rice cultivar Samba Mahsuri was identified to exhibit broad-spectrum resistance to BB. Using next-generation gene mapping approaches, we identified a genomic interval in chromosome 6 to be linked to BB resistance. Analysis of the SNPs in the locus and subsequent linkage analysis indicated that a missense SNP located in the second exon of the *Mitogen-Activated Protein Kinase 6* (*OsMAPK6*) could be the causal mutation. A detailed examination of *OsMAPK6* gene sequences in over 4000 rice genomes revealed that the mutant allele identified in this study is novel to rice germplasm. The candidate SNP causes the substitution of an invariant Serine residue with a Proline residue (S84P), thus likely affecting the protein function. Global transcriptome, metabolome, biochemical, and molecular analyses suggested that BB42 exhibit constitutive immunity. On the mechanistic front, the mutation seems to result in diminished brassinosteroid signaling in BB42. Taken together, we report a novel allele of a highly conserved *MAPK* gene and provide evidence for its possible association with constitutive immunity and disease resistance in rice.

## INTRODUCTION

Plants possess a potent innate immunity which every cell that faces a threat can elicit. The two-tiered innate immunity of plants consists of Pattern-Triggered Immunity (PTI) and Effector-Triggered Immunity (ETI) (Jones and Dangl, 2006). PTI is elicited by the recognition of the Pathogen-Associated Molecular Patterns (PAMPs) or Damage-Associated Molecular Patterns (DAMPs). PAMPs include conserved microbial-derived molecules like flagellin, peptidoglycan, LPS, and chitin; DAMPs include damage-associated molecules like extracellular ATP, extracellular DNA, and oligogalacturonides (Chen and Ronald, 2011; Pontiggia *et al*., 2020). The patterns are perceived by the membrane-bound receptor kinases, that are collectively called as Pattern Recognition Receptors (PRRs), that relay signals to the signaling proteins like the MITOGEN-ACTIVATED PROTEIN KINASE (MAPK), that typically transduce the signals to transcription factors. PTI induces an extensive transcriptional and translational reprogramming that results in the production of defense-related proteins, phytohormones, and metabolites, thus limiting pathogen entry and proliferation (D Couto, 2016; Yuan *et al*., 2021; Zipfel and Felix, 2005). On the other hand, ETI is a stronger defense response that is activated by the recognition of pathogen-secreted proteins (also known as effectors) by intracellular receptors mainly belonging to the Nucleotide-Binding and Leucine-rich Repeat containing Receptors (NB-LRRs/NLRs) superfamily. ETI is manifested by the Hypersensitive Response (HR) which results in a localized cell-death, thus restricting the spread of pathogens (Jones *et al*., 2016). On the other hand, uncontrolled regulation of various arms of the plant immune system results in constitutive immunity which in turn results in various traits like stunted growth, spontaneous lesions, and disease resistance among others (Freh *et al*., 2022).

Bacterial Blight (BB) is the most serious bacterial disease of rice that can result in up to 70% yield loss depending on the infection severity (Goto, 1992; K. Reddy, 1979; Ou, 1973, 1985). Rice cultivation in India is affected by BB with ∼8.5% loss in annual yield (Savary *et al*., 2019). BB is caused by the Gram-negative bacterium *Xanthomonas oryzae* pv. *oryzae* (*Xoo*). *Xoo* invades rice leaves through hydathodes and colonizes the extracellular spaces in xylem parenchyma tissue (Hilaire *et al*., 2007). Under high humidity conditions, the bacteria ooze out of the openings such as hydathodes or wounds and form beads or strands of exudates that potentially act as inoculum for further infections (Mew, 1993). Being a prolific colonizer of the vascular tissue, *Xoo* spreads systemically along the veins and results initially in chlorotic yellow and later in necrotic lesions on the infected leaves, ultimately leading to drying of the leaves (Joshi *et al*., 2020; Niño-Liu *et al*., 2006).

Previous studies have associated several genes and their alleles with BB resistance, of which a few alleles are widely used in breeding programme to develop BB resistant rice varieties (Fiyaz *et al*., 2022; Jamaloddin *et al*., 2021; Jiang *et al*., 2020). Such resistant varieties have been deployed in the field with ample success (Dasari *et al*., 2022; Huang *et al*., 1997; Jamaloddin *et al*., 2020; Oliva *et al*., 2019; Pradhan *et al*., 2015; Sundaram *et al*., 2008). It was observed that a combination of two to three *R*-genes was effective against majority of the *Xoo* pathotypes. However, a constant selection pressure on pathogens combined with their rapid generation time is expected to result in the evolution of adapted pathotypes that breakdown existing sources of disease resistance. Studies have identified multiple strains of *Xoo* that breakdown individual resistance genes (Kaur *et al*., 2020; Mishra *et al*., 2013; Quibod *et al*., 2020; Yugander *et al*., 2017, 2022). Adding another layer to this serious issue is the migration of hypervirulent strains from one country to another (Carpenter *et al*., 2020). Despite the existence of several *R*-genes, the practical deployment of the genes needs to be rigorously tested on a case-to-case basis to ensure the compatibility of the genes to different agroclimatic zones. For instance, although *Xa7* was identified to be a broad-spectrum R-gene, a study in India found that 94% of the 71 strains tested were virulent on a *Xa7*-positive rice line (Yugander *et al*., 2022). Therefore, to counter resistance gene breakdown and to curb disease outbreaks, it is essential to identify new sources of broad-spectrum resistance that are suitable for given agroclimatic conditions and the cultivar background.

Mutation breeding is an efficient method to obtain novel sources of resistance in relatively less time, while preserving the genetic architecture of the desired cultivar. Further, the availability of advanced gene mapping approaches enables forward genetics using populations derived from the induced mutant lines (Abe *et al*., 2012; Takagi *et al*., 2013). Earlier, we reported the generation and screening of a large-scale ethyl methanesuphonate-induced mutant lines of the rice mega-variety Samba Mahsuri for various traits of economic importance (Potupureddi *et al*., 2021). Here, we report the characterization of one of the mutant lines (henceforth BB42) that exhibited a broad-spectrum resistance to various hypervirulent isolates of *Xoo*. Through a genetic mapping study, we observed that resistance in BB42 is controlled by a single recessive locus. We further identified a SNP located in the second exon of *OsMAPK6* gene that causes Serine to Proline substitution at the 84^th^ position and cosegregates with the resistance trait. Notably, the mutant allele identified here represents a novel genetic variation in the rice germplasm. Transcriptome profiling in combination with multiple biochemical and molecular assays revealed a constitutively active basal defense signaling in BB42 when compared to SM. Particularly, several kinase-encoding genes were significantly upregulated, and several members of the major transcription factor families were differentially expressed. Overall, our study reports a novel allele of *OsMAPK6* that is linked to bacterial blight resistance, and constitutive immunity and presents a potential genetic source for breeding BB resistance in rice.

## RESULTS

### BB42 exhibits broad-spectrum resistance to highly virulent *Xoo* isolates

Earlier, we identified BB42 from an EMS-induced mutant library of SM as a line exhibiting enhanced resistance to the bacterial blight disease of rice (Potupureddi *et al*., 2021). Here, we characterized BB42 further through a combination of phenotyping, genomic, molecular, transcriptomics and metabolomics approaches. Morphologically, BB42 exhibited reduced height compared to SM (Fig. 1A, B). Further, to understand the spectrum of resistance, we challenged BB42 with several hypervirulent strains of *Xoo* that were shown to breakdown resistance provided by major resistance genes including *xa5*, *xa13* and *Xa21* (Mishra *et al*., 2013). The results revealed that BB42 exhibits enhanced resistance to all the tested *Xoo* strains (Fig. 1C, D). Notably, challenging SM and BB42 with *Xanthomonas oryzae* pv. *oryzicola* (*Xoc*) – a pathovar of *X. oryzae* that causes bacterial leaf streak (BLS) in rice - showed no difference in the susceptibility. Intriguingly, the susceptibility level of SM to *Xoc* was significantly lesser than that of Nipponbare - another BLS susceptible rice variety (Figure S1A). Taken together, these results suggest that BB42, a semi-dwarf mutant of SM, is resistant to *Xoo* in what appears to be a non-race-specific manner.

**Figure 1:**
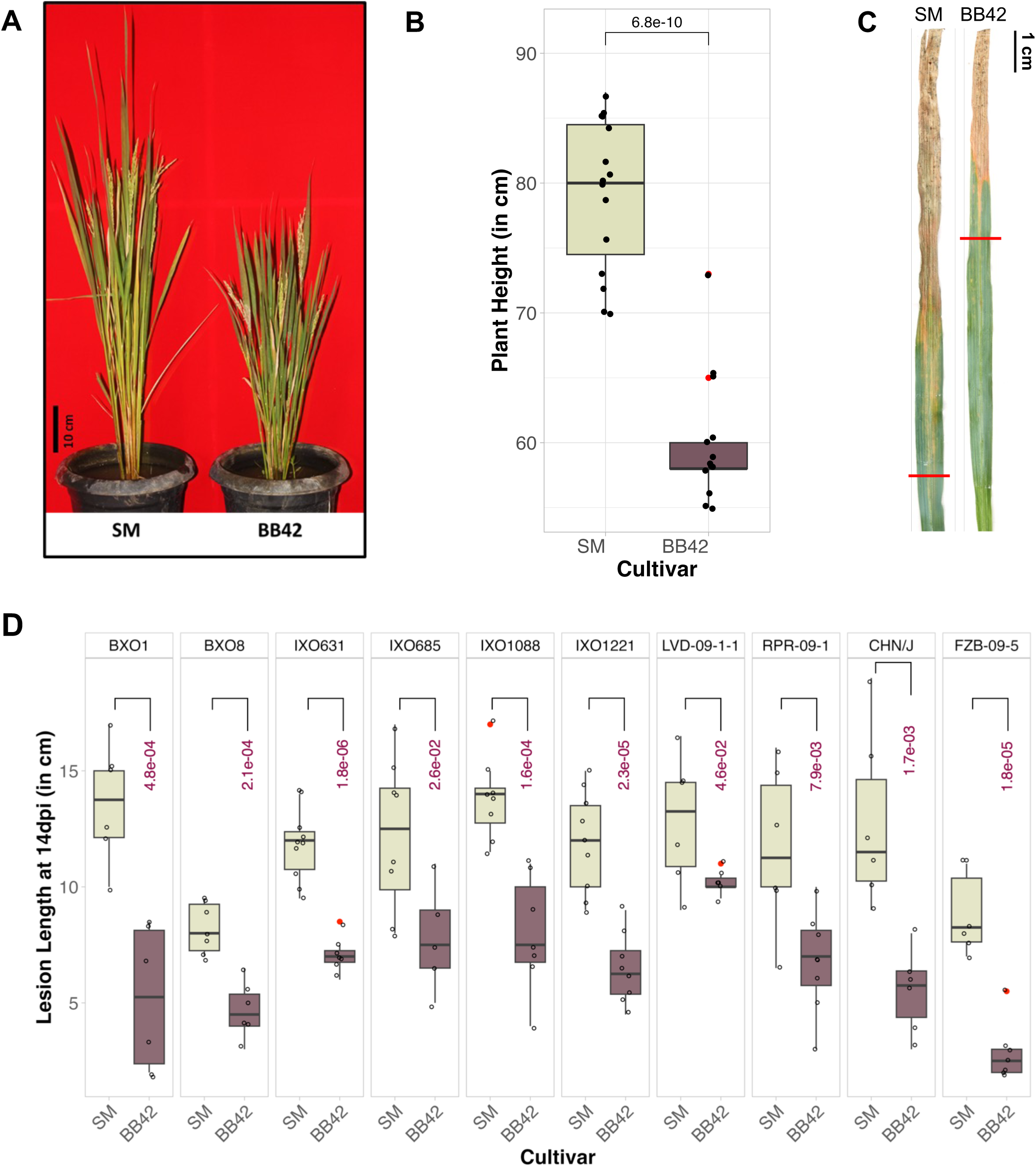
BB42 is a dwarf and bacterial blight resistant derivative of SM. (A) and (B) Comparison of the morphological features of SM and BB42 showing a significantly reduced height of the latter. The statistical significance was calculated using unpaired two-tailed Student’s *t* test. Number of samples: SM=15, BB42=13. (C) Representative leaves of SM and BB42 showing the disease symptoms at 14 days post inoculation (dpi). (D) Lesion length readings of *Xoo* infected SM and BB42 plants at 14 days post inoculation. Fully expanded leaves were clip inoculated with *Xoo* suspension at OD_600_ 1.0 (n>5). Statistical significance was calculated using one-way Anova and comparisons between groups were performed using Tukey’s HSD (Honest Significant Difference) test with a 95% family-wise confidence level. *P*<0.05 was considered statistically significant. In (B) and (D) boxes represent the 25^th^ to 75^th^ percentile while the line within the box indicates the median. The whiskers extend to maximum and minimum data points that are within 1.5 times the interquartile range. Red dots indicate outliers.

### A missense mutation in *OsMAPK6* is linked to bacterial blight resistance

We next aimed to identify the genetic basis of bacterial blight resistance in BB42. To this end, we generated two F_2_ populations, each with 244 plants, and screened them with a virulent *Xoo* strain, IXO1221. Phenotype segregation in both the populations followed the Mendelian ratio for a monogenic recessive trait (Table S4; Figure S2A, B). Progenies from both the populations were used for gene mapping through the QTL-sequencing pipeline (Takagi *et al*., 2013). The analysis revealed a genomic interval in the short arm of chromosome 6, in both the populations, to be linked to resistance (Fig. 2A, B). We identified five Single Nucleotide Polymorphisms (SNPs) within the interval, of which, one SNP was annotated as a ‘missense and splice region’ variant and was predicted to be deleterious (Table 1). This SNP (T → C at SM-Chr6:2664062) is located in the second exon of the rice *Mitogen Activated Protein Kinase 6* (*OsMAPK6*) gene and results in the substitution of a Serine by Proline (Fig. 2C). Analysis of the RNA reads mapped to OsMAPK6 indicated no anomalies in the splicing of the gene between SM and BB42 (Figure S3). Multiple sequence alignment of one-hundred MAPK6 homologues across different plant species indicated a 100% conservation of the affected Serine residue (Fig. 2D). Notably, when searched in the RiceVarMap V2.0 database (Zhao *et al*., 2021) which contains sequences of about 4500 rice accessions, we found no accession to carry any variation at this position. To further ascertain the co-segregation of the observed SNP with BB resistance, we developed PCR-based markers that differentiate the SM and the BB42 alleles (Figure S2C). We tested randomly selected F_2_ progenies that were resistant or susceptible to *Xoo* IXO1221 and found that the BB42 allele of *OsMAPK6* co-segregated with the resistant progenies, whereas the susceptible progenies were homozygous for SM allele or were heterozygous (Fig. 2E). Overall, these results suggest that the variation in the *OsMAPK6* gene likely confers BB resistance. The identified allele is novel to rice germplasm and the affected Serine residue is evidently invariable in MAPK6 homologues that were assessed, indicating a likely role of this novel variant in BB resistance.

**Figure 2:**
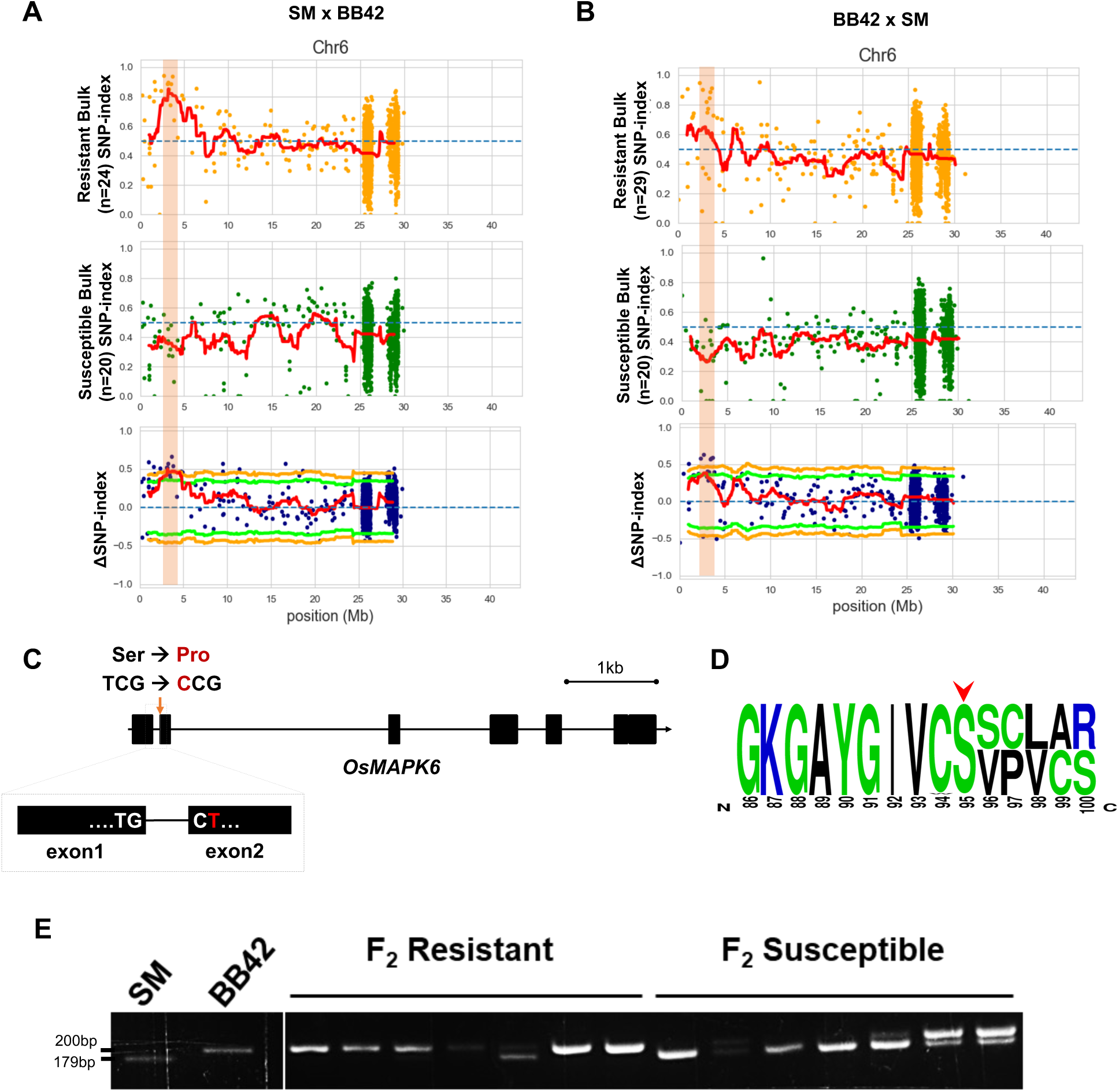
A novel mutation in *OsMAPK6* is linked to BB resistance. Allele frequency plots of chromosome 6 indicating the mapped regions in two F_2_ population: (A) SM x BB42 and (B) BB42 x SM. The top plot represents the regression analysis of SNP-index of resistant bulk, middle plot represents that of susceptible bulk, and the bottom plot indicates the delta SNP-index plot. The red line is the moving average line plotted by averaging the SNP or delta SNP indices in 2Mb windows and a 100kb step size. Each dot represents an SNP. The transparent orange bar indicates the mapped genomic interval. (C) Schematic gene diagram of *OsMAPK6* indicating the affected base (red arrow) and the amino acid substitution caused. (D) Multiple sequence alignment logo of 100 plant homologues of MAPK6 spanning the Serine residue of interest (red arrowhead). (E) Genotyping of selected F_2_ progenies using MAPK6 allele-specific PCR markers showing co-segregation of the mutant allele with resistance.

**Table 1:** List of high-frequency SNPs identified within the mapped genomic interval, their SNP indices, and effect.

| Chr | Pos<sup>a</sup> | SNP index<br>(SM x BB42) | SNP index<br>(BB42 x SM) | Effect |
| --- | --- | --- | --- | --- |
| Chr6 | 2243785 | 0.76 | 0.72 | Intron variant |
| Chr6 | 2358077 | 0.81 | 1.00 | Intron variant |
| Chr6 | 2379923 | 0.84 | 0.94 | Downstream gene variant |
| **Chr6** | **2664062** | **0.94** | **0.90** | **Missense variant & splice region variant;**<br>**SIFT - deleterious (0)** |
| Chr6 | 3057969 | 0.88 | 0.81 | Downstream gene variant |

<sup>a</sup> Coordinates in SM reference genome

### *Xoo* induces an overlapping transcriptome change in SM and BB42

Given the central role of MAPK6 in orchestrating cellular signalling during defense and development, we reasoned that BB42 and SM are likely to have different transcriptional profiles which might explain the observed resistance to *Xoo*. Therefore, we performed a global transcriptome profiling of leaves from SM and BB42 that were sampled immediately post infection (Mock) or at 24 hours post infection with *Xoo* IXO1221 (*Xoo*). The samples were analysed to identify the treatment-based transcriptome changes in both the genotypes (i.e., *Xoo* vs Mock in SM and BB42) as well as genotype-specific transcriptome differences in both the treatments (i.e., BB42 vs SM in Mock and *Xoo*). Four combinations of samples were analysed in total and the results indicated that *Xoo* treatment caused a major transcriptome reprogramming in both the genotypes (1974 DEGs in SM and 1477 DEGs in BB42) (Data S1). On the other hand, the genotype-specific differences in the transcriptome were relatively smaller (810 DEGs in Mock treatment and 493 DEGs in *Xoo* treatment) (Fig. 3A; Figure S4; Table S5). Further comparative analyses revealed that about 60% of the upregulated genes in SM and BB42 upon *Xoo* infection were common (Fig. 3B). With regards to the downregulated genes, about 87% of the genes downregulated in BB42 were also downregulated in SM upon *Xoo* infection (Fig. 3B), while several additional genes were uniquely downregulated in SM. This is suggestive of a largely similar *Xoo*-mediated transcriptional landscape in SM and BB42 at gene level, which is also reflected at pathway levels (Figure S5).

**Figure 3:**
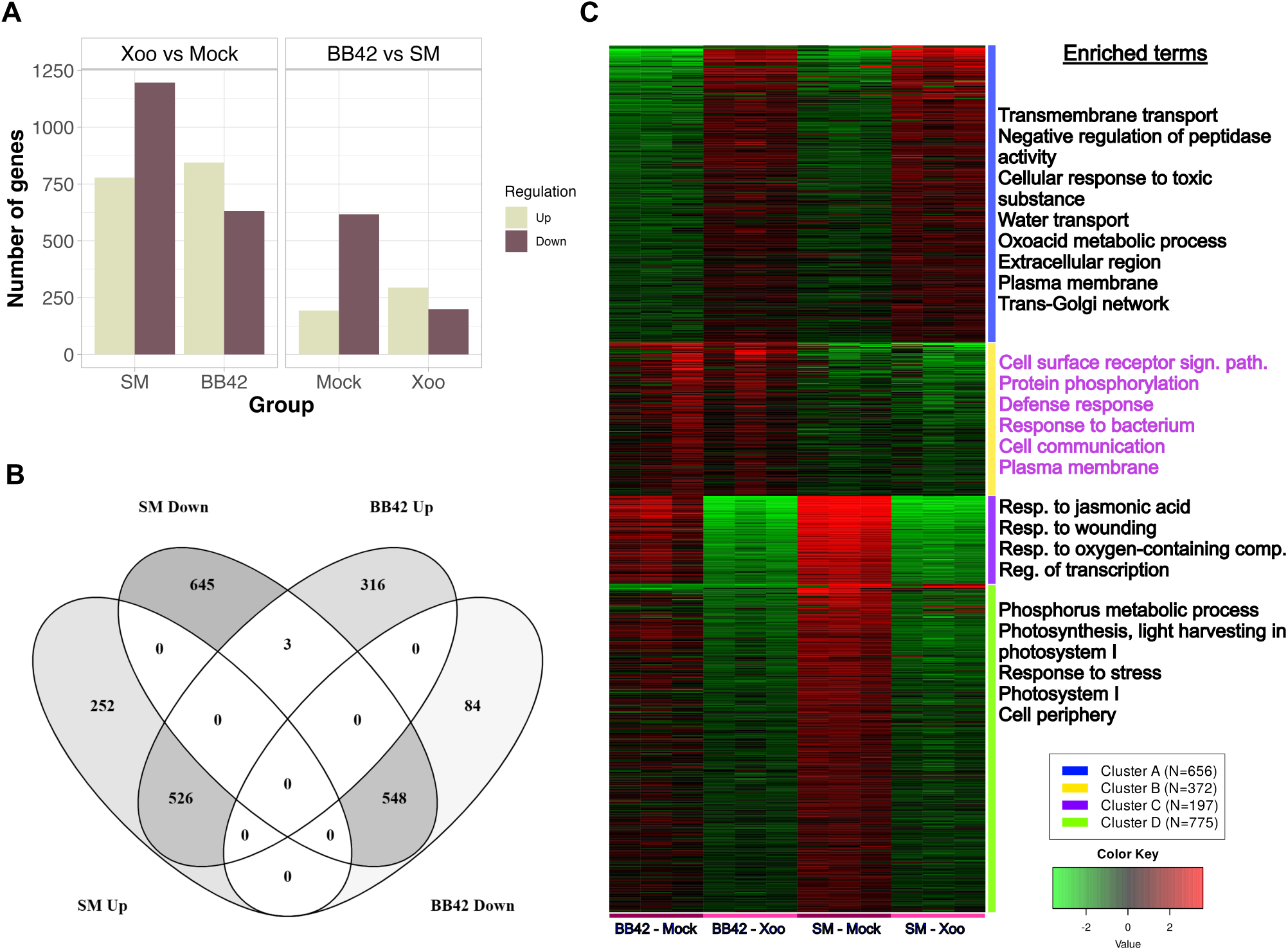
SM and BB42 share a substantial *Xoo*-mediated transcriptional landscape. (A) Number of Differentially Expressed Genes (DEGs) identified in various comparisons of the transcriptome data. (B) Venn diagram showing common and unique differential gene expression patterns in SM and BB42 samples challenged with *Xoo*. Up and Down mean upregulation or downregulation of genes in *Xoo* treated condition as compared to mock treated condition in the mentioned genotypes. (C) k-means clustering of gene expression profiles of the top 2000 genes. Lines with different colours indicate different clusters based on the gene expression values. Number of genes within each cluster is provided in the legend. Enriched terms adjacent to each cluster are the GO terms that are enriched within corresponding clusters.

We then performed a *k*-means clustering analysis of the top 2000 genes exhibiting highly variable expression across different samples and treatments. We observed 1628 genes showing similar expression trend in both SM and BB42, whereas 372 genes showed an increased expression in BB42 irrespective of the treatment, when compared to SM (Fig. 3C; Cluster B). Intrigued by this observation, we did a cluster-wise GO term enrichment analysis and noted that cluster B was enriched with genes that are involved in defense response and cell signalling. It is mention-worthy that the higher expression of defense genes is observed even in control BB42 plants (Fig. 3C), which points towards the likelihood of a constitutively active defense in BB42. Therefore, we probed deeper to understand the nature of constitutive defense in BB42 using DEGs from the mock-treated samples.

### BB42 exhibits constitutive immunity

We considered the DEGs between mock-treated BB42 and SM for further analysis to understand the basal genotype-specific differences. A GO term enrichment analysis revealed higher expression levels of genes involved in defense response, receptor signalling, and protein phosphorylation (Fig. 4A). Conversely, certain genes involved in wounding response, response to jasmonic acid, and regulation of transcription were expressed at lower levels in BB42 in comparison to SM (Fig. 4A). We first looked into the genes present in the highly expressed categories and noted the presence of various classes of defense related genes (n=41) encoding receptor kinases, receptor-like cytoplasmic kinases, a catalase, pathogenesis-related proteins and a JA biosynthetic protein (Table S6). Very importantly, amongst the above set of genes were fourteen genes encoding Nucleotide-Binding adaptor shared by APAF-1, R proteins, and CED-4 (NB-ARC) domain containing proteins and Nucleotide-Binding Leucine-rich Repeat containing Receptor proteins (NB-LRR or NLRs), which is one of the characteristic traits observed in autoimmune mutants in plants (Freh *et al*., 2022) (Figure S6). While this suggests an enhanced defense signalling in BB42, we were puzzled to observe downregulation of JA responsive genes. However, a detailed inspection indicated that such genes encoded the JASMONATE-ZIM DOMAIN (JAZ or TIFY) proteins, which act as repressors of JA signalling (Table S7). This further suggests the possibility of an active and/or enhanced JA signalling in BB42. To validate this observation and also to check if this scenario persists through different developmental stages, we performed quantitative PCR using defense marker genes (PR genes) in SM and BB42 seedlings. In accordance with the previous observation, we found significant upregulation of defense marker genes in BB42 seedlings (Figure S76A). Moreover, we found other hallmarks of defense activation like H_2_O_2_ accumulation and MAPK activation to be enhanced in BB42 even in unelicited conditions (Figure S7B, C). Further, as we observed an upregulation of various types of kinases, we used Phospho-Serine and Phospho-Threonine/Tyrosine antibodies to assess the qualitative differences in the phosphorylated proteins in SM and BB42. There was an observable difference in the signal patterns in all forms of phosphorylated proteins between SM and BB42 (Figure S7D). By and large, the transcriptomics data in combination with other assays and the dwarf stature of BB42 are indicative of constitutive immunity and a differential phosphorylation-mediated signalling between the two lines.

**Figure 4:**
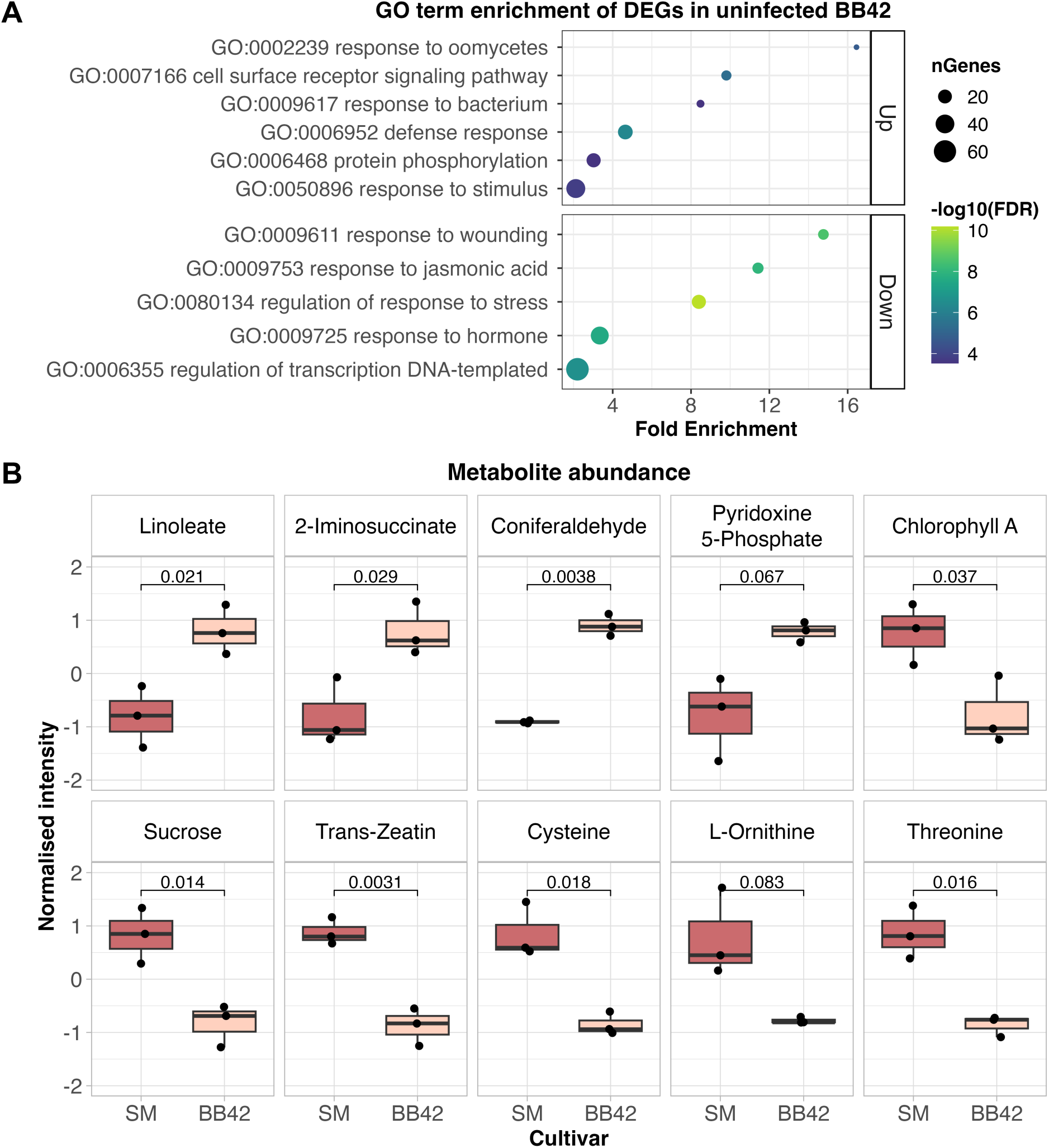
BB42 exhibits constitutive immunity. (A) Gene ontology term enrichment of the genes that are upregulated (top panel) or downregulated (bottom panel) in mock samples of BB42 as compared to the corresponding SM samples. (B) Box plots showing the abundance of certain primary and secondary metabolites that are differentially accumulated between SM and BB42. The raw peak intensities were normalised and plotted on the y-axis. The numbers above the horizontal lines indicate the *P*-values as calculated using unpaired Student’s *t*-test considering equal variance between samples. *P*<0.05 is considered statistically significant. Boxes represent the 25^th^ to 75^th^ percentile while the line within the box indicates the median. The whiskers extend to maximum and minimum data points.

We performed an untargeted metabolite profiling of SM and BB42 leaves at the tillering stage and observed a general difference in the metabolite profiles of SM and BB42 (Figure S8A). Among the detected metabolites, 92 metabolites were differentially accumulated (DAMs), of which 40 were relatively abundant in BB42 and 52 were relatively abundant in SM (Figure S8B; Data S2). Enrichment analysis of the DAMs indicated linoleic acid metabolism, vitamin B6 metabolism, and nicotinate metabolism to be upregulated and starch and sucrose metabolism, and glutathione metabolism to be downregulated in BB42 (Figure S9). On that note, levels of some of the compounds associated with the above pathways showed clear difference between SM and BB42. Notably, compounds that are associated with defense like linoleic acid and coniferaldehyde were abundant in BB42, while compounds that were associated with disease susceptibility like sucrose, *trans*-Zeatin, and amino acids were abundant in SM (Fig. 4B).

### Brassinosteroid signalling is affected in BB42

Earlier, Liu *et al*., (2015) identified an allele of *OsMAPK6* that resulted in dwarf stature and small grain size (*dsg1*) and showed that the observed traits were due to imbalance in brassinosteroid (BR) homeostasis. Interestingly, analysis of various morphological traits showed slight similarity between *dsg1* and BB42. For instance, BB42 grains were slightly yet significantly shorter than those of SM, but there were no difference in grain width (Fig. 5A, B, C). We then tested the BR sensitivity of SM and BB42 plants by measuring coleoptile elongation following treatment with 24-epibrassinolide (epiBL). The elongation of BB42 coleoptile in response to epiBL was not as significant as that of SM, suggesting a reduced sensitivity to BR in BB42 (Fig. 5D). Intriguingly, brassinolide was detected in the metabolome profiling and its level was slightly, albeit not significantly, higher in BB42 than in SM (Fig. 5E). It is possible that the reduced sensitivity to BR in BB42 might have negatively impacted the feedback inhibition of BR biosynthesis, thus resulting in higher basal BR levels. Additionally, expression of several genes involved in BR signalling and response was diminished in BB42 as compared to SM (Table S8). On the whole, it seems like BB42 is impaired in BR sensitivity and a role of the mutated OsMAPK6 in this impairment could be speculated.

**Figure 5:**
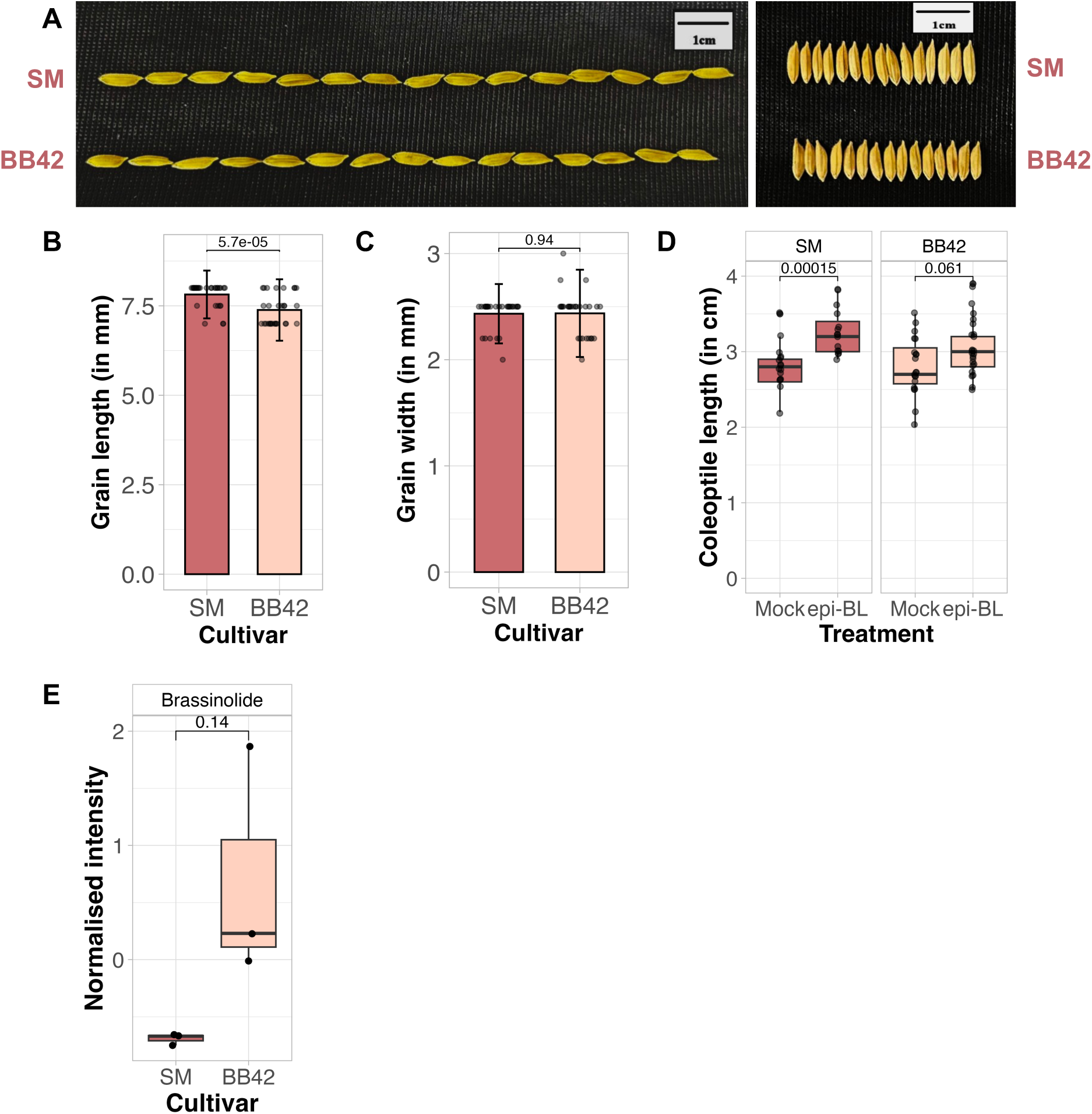
Brassinosteroid signalling is impaired in BB42. (A) Comparison of grain size of SM and BB42. Scale bar = 1 cm. (B) and (C) bar graphs depicting the grain length and grain width, respectively, of SM and BB42. N=30. Bars are drawn to average and the error bars indicate standard deviation. The dots represent the data points. (D) Box plots showing the length of the hypocotyl upon treatment of the seeds with mock solution or solution containing 5μM 24-epibrassinolide. Boxes represent the 25^th^ to 75^th^ percentile while the line within the box indicates the median. The whiskers extend to maximum and minimum data points that are within 1.5 times the interquartile range. Solid black dots outside the whiskers, if any, indicate outliers. N>10. The numbers above the horizontal line indicates *P*-values as calculated using unpaired Student’s *t*-test considering equal sample variance. *P*<0.05 is considered statistically significant. (E) Box plot showing the abundance of brassinolide in SM and BB42 leaves. Boxes represent the 25^th^ to 75^th^ percentile while the line within the box indicates the median. The whiskers extend to maximum and minimum data points.

## DISCUSSION

In this study, we report an EMS-induced allele of *Mitogen-Activated Protein Kinase 6* and show its involvement in resistance to bacterial blight, which is a serious disease of rice. The variant of OsMAPK6 identified here is new to rice germplasm wherein an otherwise invariant Serine at the 84^th^ position is replaced by Proline. Not just in plants, the affected Serine was observed to be invariant in MAPK6 homologues of different kingdoms (Figure S10A, B). Further, the serine residue is conserved in ten of the seventeen rice MAPKs (Figure S11A, B). These results signify the importance of the Serine residue for the function of MAPK6 homologues and perhaps in majority of the rice MAPKs as well. The causal SNP is a T to C transition, a non-canonical type of EMS-induced variation. EMS is believed to predominantly cause G/C to A/T type transitions, although exceptions are reported (Abe *et al*., 2012; Greene *et al*., 2003; Jhingan *et al*., 2023; Loveless, 1969; Till *et al*., 2007).

### Effect of the variation on MAPK6 function

MAPK6 homologues are highly conserved across eukaryotes and play vital roles in disease and development (Meng and Zhang, 2013; Sun and Zhang, 2022). Previous studies have associated OsMAPK6 with various growth processes (Liu *et al*., 2015; Xu *et al*., 2018; Yi *et al*., 2016) as well as defense response and resistance to diseases like blast, blight and bacterial leaf streak (Chen *et al*., 2021; Chujo *et al*., 2014; Kim *et al*., 2012; Kishi-Kaboshi *et al*., 2010; Lieberherr *et al*., 2005; Xu *et al*., 2018). It is interesting to note that natural variations in *OsMAPK6* differentially affect the protein activity and resistance to *Xoc* (Ma *et al*., 2021). In line with this, the observed difference between SM and Nipponbare in their response to *Xoc* (Figure S1A) might be explained by the natural variation in the N-terminus of MAPK6 between SM and Nipponbare (Figure S1B). Further, certain loss-of-function alleles of *OsMAPK6* were reported to cause embryo lethality (Ma *et al*., 2021; Yi *et al*., 2016). It was also observed that a heterozygously inherited loss-of-function mutation in OsMAPK6 could confer moderate resistance to *Xoo* and *Xoc* (Ma *et al*., 2021). Hence, the effect of the S84P mutation on OsMAPK6 function becomes a primary question. The fact that the activation of OsMAPK6 is not affected in BB42 (Figure S6C) suggests that the mutation has not abolished the protein activation and function, rather the protein’s function could have been altered. Intriguingly, one of resistant progenies showed a heterozygous genotype in our analysis. Noteworthy, the SNP-index of this variation is not 1.0 in either of the population. This might indicate an effect of the environment, that could have resulted in one or a few heterozygous individuals showing resistance trait getting included in our sequencing and downstream analyses. Nevertheless, none of the susceptible individuals tested were homozygous for the mutant allele and all of the lines that were homozygous for the mutant allele were resistant. On balance, it is possible that the presence of the mutant allele and allele dosage in combination with the environment might contribute to BB resistance.

### MAPK6 integrity could be crucial to prevent constitutive immunity in rice

The phenotypes observed in BB42 including dwarf stature and constitutive immunity, seem to partly resemble a defective MAPK cascade reported in the model plant *Arabidopsis thaliana*. In Arabidopsis, three major MAPKs that are involved in plant immunity and pathogen response are AtMPK3, AtMPK4, and AtMPK6 (Sun and Zhang, 2022). Several studies have elucidated the signalling cascade involving AtMPK4 and showed that the MEKK1-MKK1/2/6-MPK4 cascade is involved in the negative regulation of innate immunity (Gao *et al*., 2008; Lian *et al*., 2018; Rodriguez *et al*., 2010). The mutants *mekk1, mkk1/2,* and *mpk4* exhibit stunted growth, severe dwarfism, and enhanced basal defense responses. Subsequent research showed that the MPK4 cascade is guarded by NLR proteins SUMM2 and RPS6, which get activated when the cascade is disrupted (Lian *et al*., 2018; Zhang *et al*., 2012, 2017). In this context, it is possible that S84P mutation in OsMAPK6 may modify its interactome, which in turn might activate the immune components guarding it.

As activation of MAPKs is critical for proper elicitation of plant immunity, the observation of BB42 exhibiting constitutive defense traits which includes MAPK6 activation raises the following question: Is OsMAPK6 activation in unelicited BB42 a cause or consequence of the active defense? A remarkable phenomenon observed in BB42 was the basal upregulation of several NLR-encoding genes, which appears to phenocopy the Arabidopsis plants that express a constitutively active form of MPK3 (CA-MPK3). Studies have revealed that expression of CA-MPK3 causes enhanced expression of NLR encoding genes in a salicylic acid-dependent manner (Genot *et al*., 2017; Lang *et al*., 2017, 2022). Expression of CA-MPK3 also resulted in resistance to the bacterial pathogen *Pseudomonas syringae* (Lang *et al*., 2017). Another study showed that long-lasting activation of MPK3/MPK6 is necessary for ETI-mediated reduction in photosynthetic efficiency in Arabidopsis (Su *et al*., 2018). In line with this, we observed a reduction in Chlorophyll A in BB42 as compared to SM (Fig. 4B). Further, prolonged activation of MPK3/MPK6 was observed during ETI and such activation was critical for the regulation of SA-responsive genes (Tsuda *et al*., 2013). This study suggested that the activation duration of MAPK is a determinant of the robustness of immunity. Of note, the consequence of CA-MAPK expression is dependent on the MAPK in question. For instance, overexpression of CA-MPK4 dampened immune responses and abolished pathogen resistance in Arabidopsis (Berriri *et al*., 2012). Considering the above-discussed points, it is possible that the activation status of the OsMAPK6-S84P variant and NLR expression could be correlated. Nonetheless, the directionality of these two observations is not certain yet. Also, unlike SA in Arabidopsis, our data suggests a possible involvement of JA in the execution of MAPK6-mediated defense responses in rice. Future investigations are needed to draw a firm conclusion.

It appears that BB42 is a unique mutant which possesses most hallmarks of autoimmunity except the spontaneous lesions that occur due to localised cell death. In rice, almost all the reported autoimmune mutants display spontaneous lesions on the leaves, thereby earning the name lesion mimic mutants (LMMs); some LMMs show reduced photosynthetic capacity as well (Freh *et al*., 2022; Xiaobo *et al*., 2020). Interestingly, loss of function mutations in components of MAPK signalling including OsMKK6 and OsMAPK4 have been reported to cause spontaneous lesions and autoimmunity in rice (Jiang *et al*., 2023; Wang *et al*., 2021). Although the molecular mechanisms of autoimmunity in rice is still unclear, based on the data presented here, we propose that a module involving MAPK6, NLRs, and JA signalling proteins might govern autoimmunity.

### Potential link between MAPK6, BR, and disease resistance

The relationship between OsMAPK6 function and brassinosteroid homeostasis has been established earlier (Liu *et al*., 2015) and has also been observed in our study (Fig. 5). However, whether the perturbed BR homeostasis has any direct effect on disease resistance remains to be explored. Involvement of BRs in plant-pathogen interaction as well as growth and development has been demonstrated earlier. BRs can act both synergistically and antagonistically with the immune components to regulate growth and defense (Albrecht *et al*., 2012; Belkhadir *et al*., 2012; Heese *et al*., 2007; Liao *et al*., 2020). In rice, BRs were shown to suppress rice immunity against a root pathogen (De Vleesschauwer *et al*., 2012). Another study showed exogenous treatment of rice leaves with brassinolide enhanced resistance to *Xoo* (Nakashita *et al*., 2003). The *Xanthomonas* effector Xoo2875 was shown to suppress immunity by interacting with OsBAK1, a component of PTI and BR signalling (Yamaguchi *et al*., 2013). In Arabidopsis, BRs were shown to antagonize JA-mediated plant defense (Liao *et al*., 2020). Strikingly, several components of BR signalling were downregulated whereas JA signalling was positively regulated in BB42 (Tables S6, S7, S8; Fig. 4A). Whether and how OsMAPK6-S84P affect BR homeostasis and its relation with disease resistance warrants further investigations.

Our study leads to several questions such as: Is the novel MAPK6 variant constitutively active? Does the variation affect the interactome of OsMAPK6? Are rice MAPK cascades guarded by NLRs? Is rice NLR expression controlled by OsMAPK6-mediated signalling? How could OsMAPK6 govern BR homeostasis, and what is the role of BR signalling in rice-*Xoo* interaction? Do rice pathogens directly target the components of MAPK cascade to promote virulence? BB42 can also serve as a novel source of BB resistance as initial observations suggest that there is no drastic effect on yield. However, further studies are needed to determine if BB42 has more subtle reductions in yield as compared to the parental line. More research directed towards the above questions would provide deeper insights into rice MAPK signalling, broaden our understanding of plant immunity in crop species and also indicate whether BB42 can be a source of resistance that can be used for rice improvement.

## EXPERIMENTAL PROCEDURES

### Plant materials and bacterial strains

Plant materials used for this study include the wildtype/parental cultivar Samba Mahsuri (SM; BPT5204) and the EMS-induced mutant line named BB42. Nipponbare rice cultivar was used as a susceptibility check for bacterial leaf streak disease scoring. *Xanthomonas oryzae* pv. *oryzae* (*Xoo*) and *X. oryzae* pv. *oryzicola* (*Xoc*) isolates were cultured from the lab stocks and the stocks obtained from Indian Institute of Rice Research, Hyderabad. The bacterial strains that were used are listed in Table S1. The seeds were germinated on Petri plates at 28℃ in darkness for 72 hours and under 12-hour photoperiod from days 4 to 7. Later the seedlings were potted onto soil before transplanting to the field after 14-21 days of potting and grown in greenhouse conditions under natural lighting.

### Plant inoculation

Plants in the maximum tillering stage (50-60 days) were inoculated with *Xoo* isolates using the clip inoculation method (Kauffman *et al*., 1973). The *Xoo* isolates were grown in Peptone-Sucrose (PS) broth (1% (w/v) peptone and 1% (w/v) sucrose) for 24 hours and the cells were pelleted. The cells were further washed in sterile Milli-Q water one to two times and adjusted to an optical density at 600nm (OD_600_) of 1.0. Sterile scissors were used to dip into the bacterial suspension and clip the fully expanded leaves ∼3-4 cm from the leaf tip. The disease severity was scored by measuring the lesion length from the site of inoculation after 14 days of inoculation. The scoring was performed following the Rice Standard Evaluation System (IRRI, 2014). Inoculations were done in green house with temperature maintained under 30℃ for transcriptome analysis studies and multiple-isolate screening. The parental lines and F_2_ progenies were screened in the rice field facility of CSIR-CCMB (17.41⁰ N, 78.55⁰ E) during Kharif 2020 with the *Xoo* strain IXO1221. *Xoc* inoculation was performed in greenhouse conditions on 2-3 weeks old seedlings through syringe infiltration of *Xoc* suspension at OD_600_ = 1.0. Lesion lengths were measured at 5 days post inoculation.

### Generation of mapping populations

Two mapping populations were generated in this study. The populations were obtained by using SM as recipient and BB42 as donor (SM x BB42) and vice versa (BB42 x SM). The F_1_ plants, so generated, were tested for hybridity by Sanger sequencing of known SNPs between SM and BB42, which were identified using whole genome sequencing data. The confirmed F_1_ plants were let to self-fertilize to obtain two independent segregating populations. F_2_ populations with 244 progenies per population were screened for BB resistance as described above.

### Bulk sequencing and analysis

F_2_ plants from both the populations showing the extreme phenotypes i.e., resistance and susceptibility to BB infection were selected to constitute the resistant and susceptible bulks, respectively. An equal amount of leaves (15mg per leaf) were pooled from F_2_ plants showing identical phenotype for DNA isolation. DNA from the bulked leaves as well as the parental leaves were isolated using the CTAB method and quantified using Qubit^TM^ High-Sensitivity dsDNA Assay in a Qubit^TM^ 4 fluorometer (Thermo Fisher Scientific, USA). The quality of the DNA was assessed on 0.8% agarose gel through electrophoresis. An amount of 1µg DNA from all the samples was used to prepare sequencing libraries with TruSeq Nano DNA HT Sample Preparation Kit (Illumina, USA) and qualified libraries were sequenced on Illumina NovaSeq 6000 platform to obtain 2×150 bp reads at approximately 40X coverage. The library preparation and sequencing were performed at the NGS facility of CSIR-CCMB. The sequenced reads were quality-checked using *FastQC* and were provided as input for QTLseq pipeline (Sugihara *et al*., 2020) to identify the causal loci. SM reference genome (assembled in-house) was used for the gene mapping analysis (Rao *et al*., *in preparation*). *AUGUSTUS* in the *Galaxy* platform was used for annotating the interval of interest (Afgan *et al*., 2018; Stanke *et al*., 2004). *SnpEff* was used for annotating the variants (Cingolani *et al*., 2012).

### Development of SNP markers and linkage analysis

Two derived Cleaved Amplification Polymorphic Site (dCAPS) markers were developed to distinguish the *OsMAPK6* alleles of SM and BB42. One marker was developed by introducing a SacI restriction site in the SM allele while the other marker was developed by introducing a SmaI restriction site in the BB42 allele. The primer sequences were designed based on the SM reference sequence using dCAPS Finder 2.0 (Neff *et al*., 2002). All the reactions were performed using 2X KAPA Taq ReadyMix (Roche, Switzerland) with 500-1000 pmoles/μl primers and 20-40ng of the template. The reaction conditions are as follows: 95℃ for 5 minutes, 35 cycles of 95℃ for 30 seconds, 58℃ for 30 seconds and 72℃ for 30 seconds, and 72℃ for 5 minutes. The PCR samples were then added with 0.5-1μl of corresponding restriction enzymes and incubated at 37℃ for 6 hours to overnight before resolving them on a 2.5% agarose gel for 1 hour at 100V. Gel images were captured on a gel documentation system (AlphaInnotech, USA). Further, another marker was developed based on the tetra-primer amplification refractory mutation system (tetra-ARMS) PCR (Ye *et al*., 2001). A typical tetra-ARMS-PCR reaction contains four primers among which each of the two inner primers is specific to the two alleles of interest. The tetra-ARMS reaction to distinguish the *OsMAPK6* alleles was standardized, and the reaction composition is as follows: 500 nM of each primer, PCR buffer – 10mM tris-HCl (pH8.3), 20mM ammonium sulphate, 2mM magnesium chloride, and 0.25mM of each dNTP, 0.2μl of 5U/μl Taq DNA polymerase (G-biosciences), and 40ng of the template. The reaction conditions are as follows: 95℃ for 5 minutes, 35 cycles of 95℃ for 30 seconds, 58℃ for 20 seconds and 72℃ for 30 seconds, and 72℃ for 5 minutes. The samples were resolved on a 2.5% agarose gel for one hour at 100V. To test the linkage between the OsMAPK6 allele and BB resistance, we tested the genotype of seven resistant progenies and seven susceptible progenies from the F_2_ generation using the dCAPS marker that was developed as mentioned above.

### Protein sequence analysis

Homologous sequences of OsMAPK6 were retrieved from NCBI through a BLASTp search. Sequences of all the rice MAPKs were obtained from RGAP database (Hamilton *et al*., 2025). Multiple sequence alignment (MSA) was performed using Multalin (Corpet, 1988). ESPript 3 was used to visualize MSA results (Robert and Gouet, 2014). WebLogo was used to generate sequence alignment logos (https://weblogo.berkeley.edu/logo.cgi) (Crooks *et al*., 2004). One-click mode of Phylogeny.fr server was used for constructing the maximum likelihood trees of MAPK6 homologues and the rice MAPKs (Dereeper *et al*., 2008). Phylogenetic trees were annotated using iTOL (http://itol.embl.de/) (Letunic and Bork, 2021).

### RNA sequencing and analysis

SM and BB42 plants were inoculated with the *Xoo* isolate IXO1221 during the maximum tillering stage. One-centimeter-long leaf samples from the site of clipping were collected at the time of infection as control (0 hour/Mock) and 24-hour post infection (Xoo) from both SM and BB42. The samples were collected from three independent plants serving as biological replicates. The samples were flash-frozen immediately after collection and stored at −80℃ until further processing. RNA was isolated from the samples using RNeasy Plant RNA Isolation Kit (Qiagen, Germany) with on-column DNA digestion using RNase-Free DNase Set (Qiagen, Germany). Purified RNA was quantified using Qubit^TM^ RNA HS Assay Kit (Thermo Fisher Scientific, USA). RNA integrity was assessed using agarose gel electrophoresis and BioAnalyzer 2100 (Agilent Technologies, USA). 1 µg of intact total RNA samples were used for library preparation using Illumina TruSeq Stranded Total RNA with Ribo-Zero Plant (Illumina, USA). Sequencing was performed on Illumina’s NovaSeq6000 platform with cycling conditions to obtain 2×100bp reads on an S2 flow cell.

The raw data were obtained, and the quality was assessed using *FastQC*. Adapters and low-quality reads were trimmed using *Trim Galore! RNA-STAR* was used to align the trimmed reads to the rice genome (MSU Rice Genome Annotation Project - version 7). Reads that mapped to the genes were quantified using *featureCounts*. Differential gene expression was obtained using the R library *DEseq2.* Genes with a false discovery rate ≤ 0.05 and log_2_ fold change ≤ −1 or ≥ 1 were considered differentially expressed genes (DEGs). For gene ontology, and pathway analyses ShinyGO was used. iDEP.96 was used for *k*-means clustering and further analysis (Ge *et al*., 2018). *Venny* was used for generating Venn diagrams (Oliveros, J.C., 2007). Protein domain enrichment among the DEGs was performed in STRING v12.0 (Szklarczyk *et al*., 2023). RNA sequence statistics are provided in Table S3.

### Quantitative PCR for defense marker genes

Complementary DNA (cDNA) was synthesized from 1-5 microgram RNA using EcoDry cDNA Synthesis Kit (Takara) containing Oligo(dT). The reaction condition is as follows: 42℃ for 60 minutes; 70℃ for 10 minutes. The synthesized cDNA samples were diluted to 15-25ng/μl for quantitative PCR (qPCR). qPCR was performed using PowerSYBR kit (Invitrogen) in triplicates per primer-sample combination. The reaction composition is as follows: 5μl of 2X master mix, 0.5μl each of forward primer (5μM) and reverse primer (5μM), 1μl template, 3μl nuclease-free water. The reactions were performed in 384-well plates at the following conditions: 50℃ for 2 minutes, 95℃ for 10minutes, 35 cycles of 95℃ for 10 seconds and 60℃ for 30seconds and melt curve analysis between 60℃ and 95℃. The reactions were set in Viaa7 (Applied Biosystems) or CFX384 (BioRad) instruments. The primers for qPCR reactions were designed using QuantPrime (Arvidsson *et al*., 2008). The list of primers and corresponding sequences are provided in Table S2.

### Immunoblotting

For MAPK activation assay, one-week old SM and BB42 roots were treated with mock or 1µM flg22 for 15 minutes. For assay in seedlings, 1-week laboratory-grown seedlings were harvested and processed further. Protein samples were isolated from the above samples, resolved on a 12% polyacrylamide SDS gel, transferred onto PVDF membrane and probed using the anti-Phospho-erk1/2 antibody (#9101; 1:2000 dilution; Cell Signaling Technologies). The blots were stained with Ponceau-S-stained for loading control.

For qualitative assessment of total phosphorylated proteins, SM and BB42 leaf samples from plants in maximum tillering stage (∼60 days) were processed as mentioned above and probed using anti-Phosphoserine (ab9332; 1:1000; Abcam) and anti-Phosphothreonine/tyrosine (#9381; 1:3000; Cell Signaling Technologies) antibodies. Ponceau-S-stained blot is used as loading control blots. The immuno signals were visualized using HRP-conjugated anti-rabbit secondary antibody (ab98440; 1:20000; Abcam) and ECL Western high-sensitivity substrate (Thermo Fisher Scientific, USA).

### BR sensitivity assays

All the assays pertinent to brassinosteroid treatment were performed as described earlier (Liu *et al*., 2015). Briefly, for coleoptile elongation assay, seeds of SM and BB42 were soaked overnight in fresh water, washed and plated on Petri plates containing solvent control or 5µM 24-epibrassinolide (Real-Gene Labs, UP, India) and grown further. The hypocotyl length was measured one-week post germination in dark conditions. Grain parameters like length and width were measured using a graduated ruler.

### Metabolite profiling

Metabolite profiling and analysis was performed on field-grown SM and BB42 leaves during maximum tillering stage essentially by following the steps as mentioned elsewhere (Gokulan *et al*., 2024). Sample preparation and processing were outsourced to Novelgene Technologies Pvt. Ltd., Hyderabad, India.

### Statistical analyses

The relevant statistical tests and considerations are provided in appropriate figure legends. Box plots, bar plots, and multivariate dot plots were plotted on R4.2.2 using *ggplot* package and the statistical parameters were integrated to the plots using the *ggpubr* package.

## Author contributions

RVS, MSM, and HKP: conceptualization; RVS, MSM, HKP, and GCG: methodology; GCG: formal analysis; GCG, MJ, DR, NG, ARS, NS, and DN: investigation; RVS, HKP, ST, MSM, RMS, KMB, and GSL: resources; GCG: data curation; GCG: writing - original draft; GCG, RVS, HKP, MSM, KMB, and GSL: writing - review & editing; GCG: visualization; RVS, HKP, and MSM: supervision; RVS, HKP, and MSM: funding acquisition.

## Acknowledgements

We thank the CCMB NGS Facility for nucleic acid sequencing services. We thank the skilled workers at ICAR-IIRR and CSIR-CCMB for their support in field activities and maintenance. We thank Sebastien Santini (CNRS/AMU IGS UMR7256) and the PACA Bioinfo platform for the availability and management of the phylogeny.fr website. GCG, MJ, DR, NG, ARS, NS, and DN thank the Council of Scientific and Industrial Research for fellowship. GCG thanks the Department of Biotechnology for the MK Bhan - Young Researcher Fellowship.

## Supplementary information

Figure S1: SM and BB42 are equally susceptible to *Xoc*.

Figure S2: F_2_ phenotyping and marker development.

Figure S3: Mutation in OsMPAK6 does not affect gene splicing.

Figure S4: PCA plot of gene expression.

Figure S5: Pathway enrichment analysis among DEGs.

Figure S6: Protein domain enrichment among DEGs.

Figure S7: BB42 shows signs of autoimmunity.

Figure S8: Metabolite analysis of SM and BB42.

Figure S9: Pathway enrichment analysis of differentially accumulated metabolites.

Figure S10: Phylogeny of MAPK6 homologues across different forms of life.

Figure S11: Phylogeny of seventeen OsMAPK proteins.

Table S1: List of strains.

Table S2: List of primers.

Table S3: RNA-seq read statistics.

Table S4: Phenotype segregation analysis.

Table S5: DEG statistics.

Table S6: Differential expression of defense related genes.

Table S7: Differential expression of JAZ-encoding genes.

Table S8: Differential expression of BR-related genes.

Data S1: List of DEGs in across different comparisons.

Data S2: List of differentially accumulated metabolites.

## Data availability

The DNA sequence data of the F_2_ bulks are deposited in NCBI SRA under the BioProject ID PRJNA1235664. The RNA-seq data are submitted in NCBI Gene Expression Omnibus under the accession GSE291968. The DNA sequence data of parental lines are reported in (Potupureddi *et al*., 2021). Any other materials and data are available upon reasonable request.

