## Supplementary material for "A novel allele of *Mitogen Activated Protein Kinase 6* is linked to disease resistance and constitutive immunity in rice": Figure S

### Supplementary Figures

#### Supplementary Figure 1 to 11.

A novel allele of *Mitogen Activated Protein Kinase 6* is linked to disease resistance and constitutive immunity in rice.

Gokulan C.G.[1,6,†], Mohammed Jamaloddin[1,†], Deepti Rao[1], Namami Gaur[1,2], Amita Rani Supriyo[1], Nisha Sao[1], Deepak Niranjan[1,7], Shrish Tiwari[1], Gouri S. Laha[3], Kalyani M. Barbadikar[3], Raman Meenakshi Sundaram[3], Sheshu Madhav Maganti[3,4,*], Hitendra K. Patel[1,5,*], Ramesh V. Sonti[1,7,*]

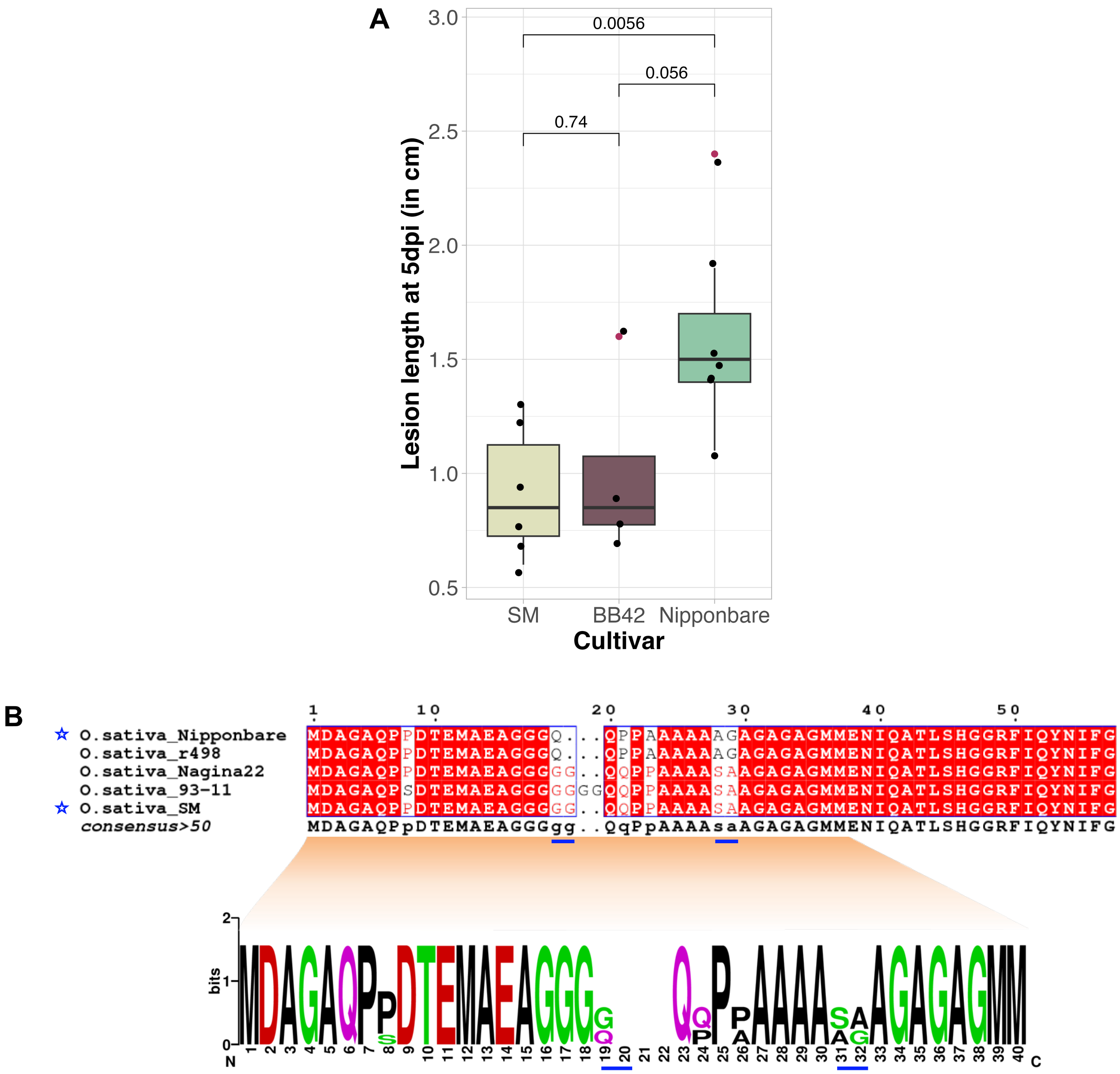


**Supplementary Figure S1:** (A) Box plots showing the lesion lengths caused by *Xanthomonas oryzae* pv. *oryzicola* (*Xoc*) on the leaves of Samba Mahsuri (SM), BB42, and Nipponbare. Bacterial suspension of the *Xoc* strain BXOR1 was adjusted to $OD_{600nm}$ 1.0 and infiltrated into the leaves of 14–21-day old seedlings. Boxes represent the 25th to 75th percentile while the line within the box indicates the median. The whiskers extend to maximum and minimum data points that are within 1.5 times the interquartile range. Red dots indicate outliers. The numbers above the horizontal lines indicate *P*-values calculated using unpaired Student's *t*-test considering equal variance. *P*<0.05 is considered statistically significant. (B) Multiple sequence alignment and sequence logo of the N-terminus of OsMAPK6 from indicated rice cultivars. The amino acid stretches with variations are underlined in blue color. Stars indicate Nipponbare and SM sequences of OsMAPK6.

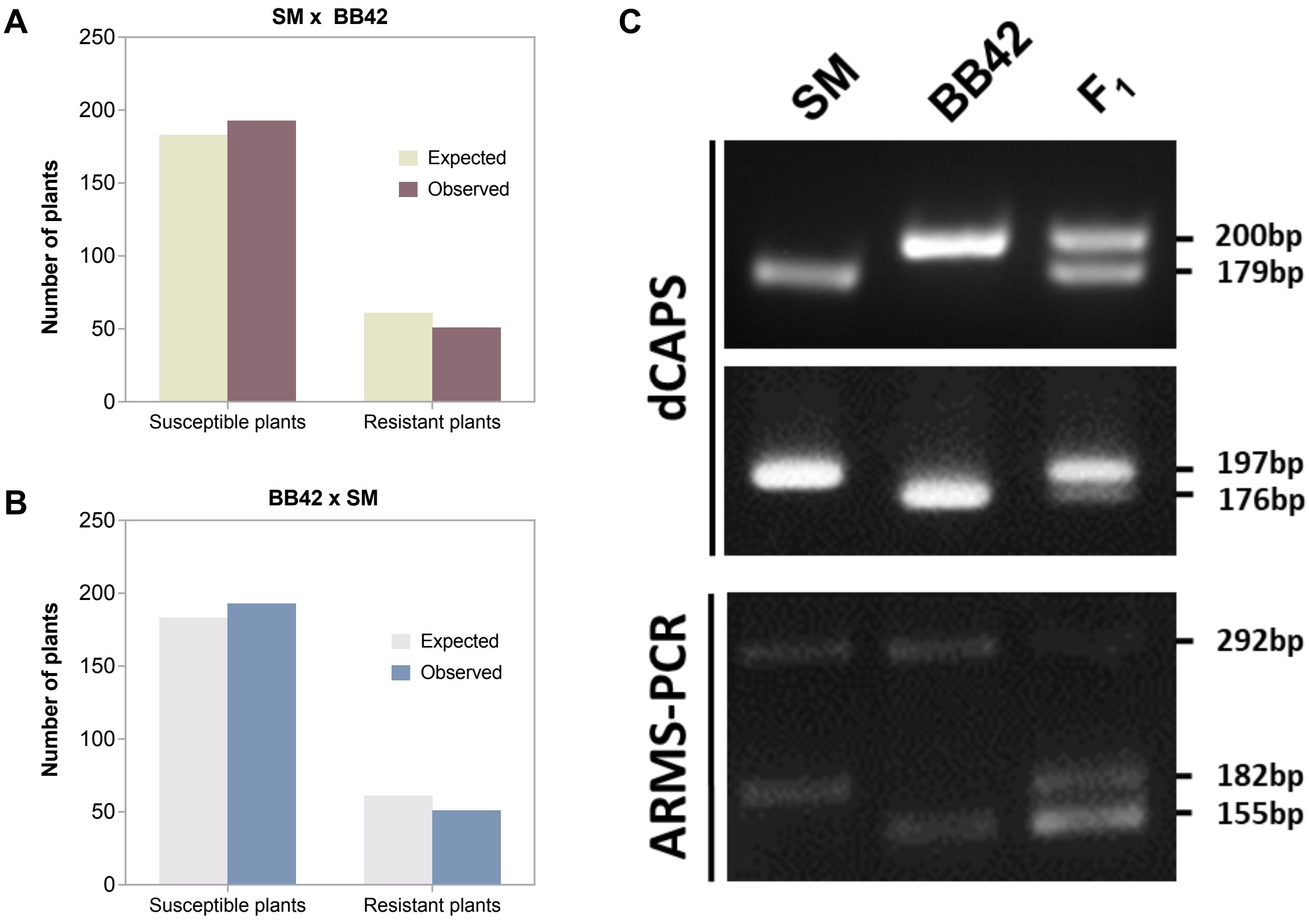


**Supplementary Figure S2:** (A) and (B) Bar plots showing the segregation of the $F_2$ populations (n=244 each) obtained from two crosses. SM x BB42: Samba Mahsuri (SM) used as recurrent parent and BB42 used as donor parent, BB42 x SM: BB42 used as recurrent parent and SM used as donor parent. (C) Band patterns of independent markers that distinguish the Samba Mahsuri (SM) and BB42 alleles of *OsMAPK6*. dCAPS: Derived Cleaved Amplified Polymorphic Sequences, ARMS-PCR: Amplification Refractory Mutation System-PCR.

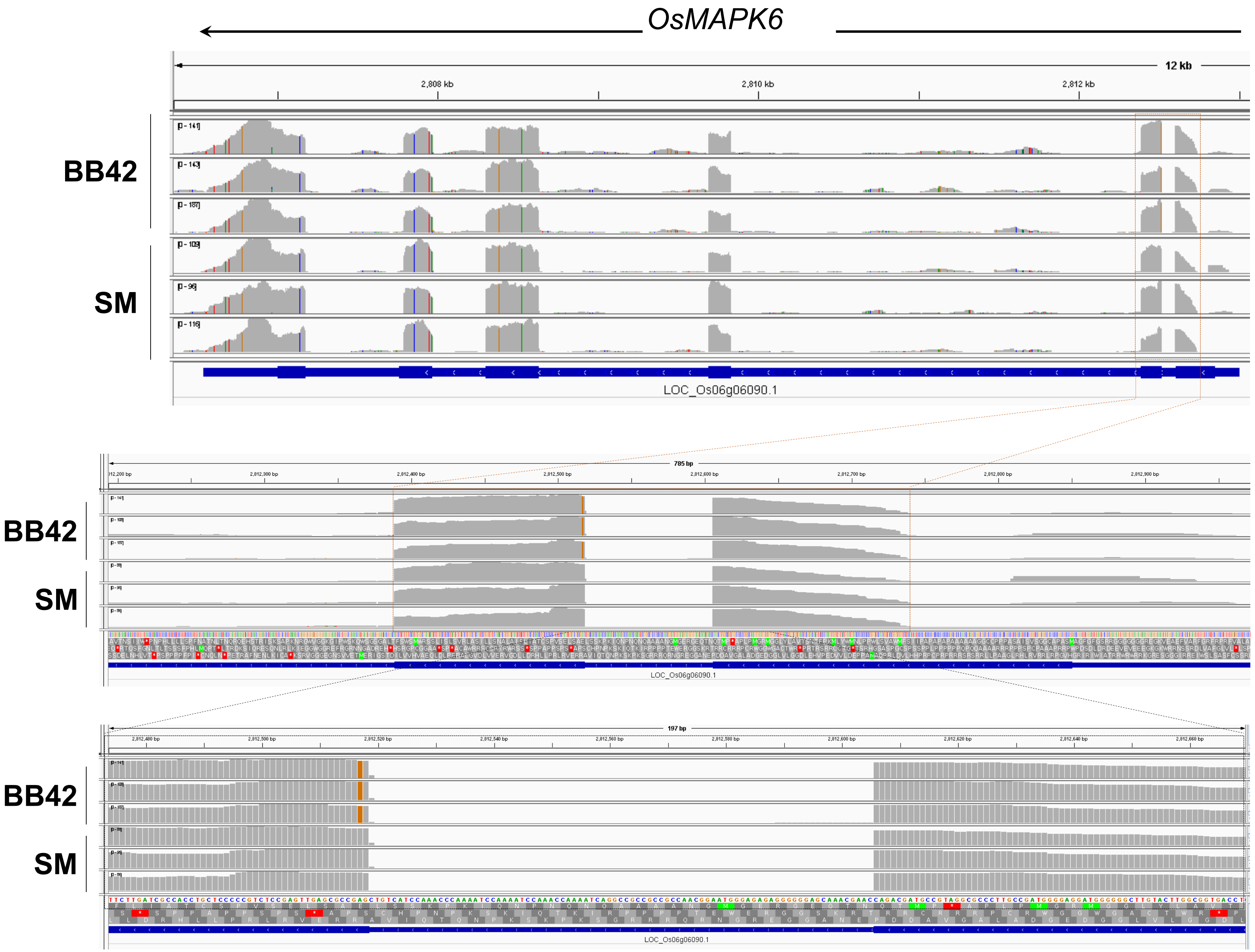

OsMAPK6
12 kb
2,808 kb
2,810 kb
2,812 kb
BB42
SM
LOC_Os06g06090.1
785 bp
BB42
SM
LOC_Os06g06090.1
197 bp
BB42
SM
LOC_Os06g06090.1

**Supplementary Figure S3:** Alignment of the RNA-sequencing reads to *OsMAPK6* using Nipponbare reference was visualized using the Integrated Genome Viewer. Top panel shows the snapshot of the entire *OsMAPK6* locus, the middle panel shows the first two exons of the gene, and the bottom panel shows a zoomed in view of the location of the SNP (brown bars in BB42) and part of the first two exons separated by an intron. The arrow on the top indicates the orientation of the gene in the genome.

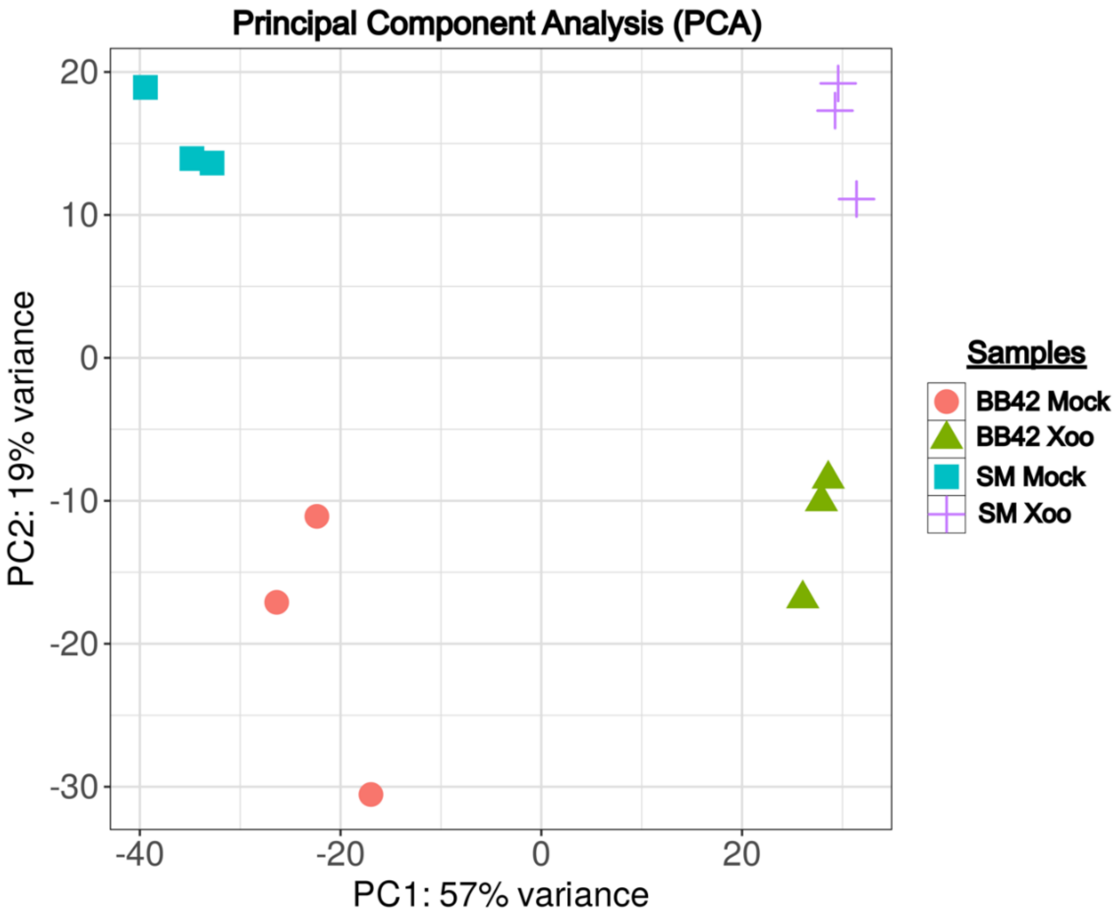


**Supplementary Figure S4:** Principal Component Analysis (PCA) plot of the Samba Mahsuri (SM) and BB42 transcriptome datasets. Mock: 0hpi experiment, *Xoo*: *Xanthomonas oryzae* pv. *oryzae* indicates 24hpi

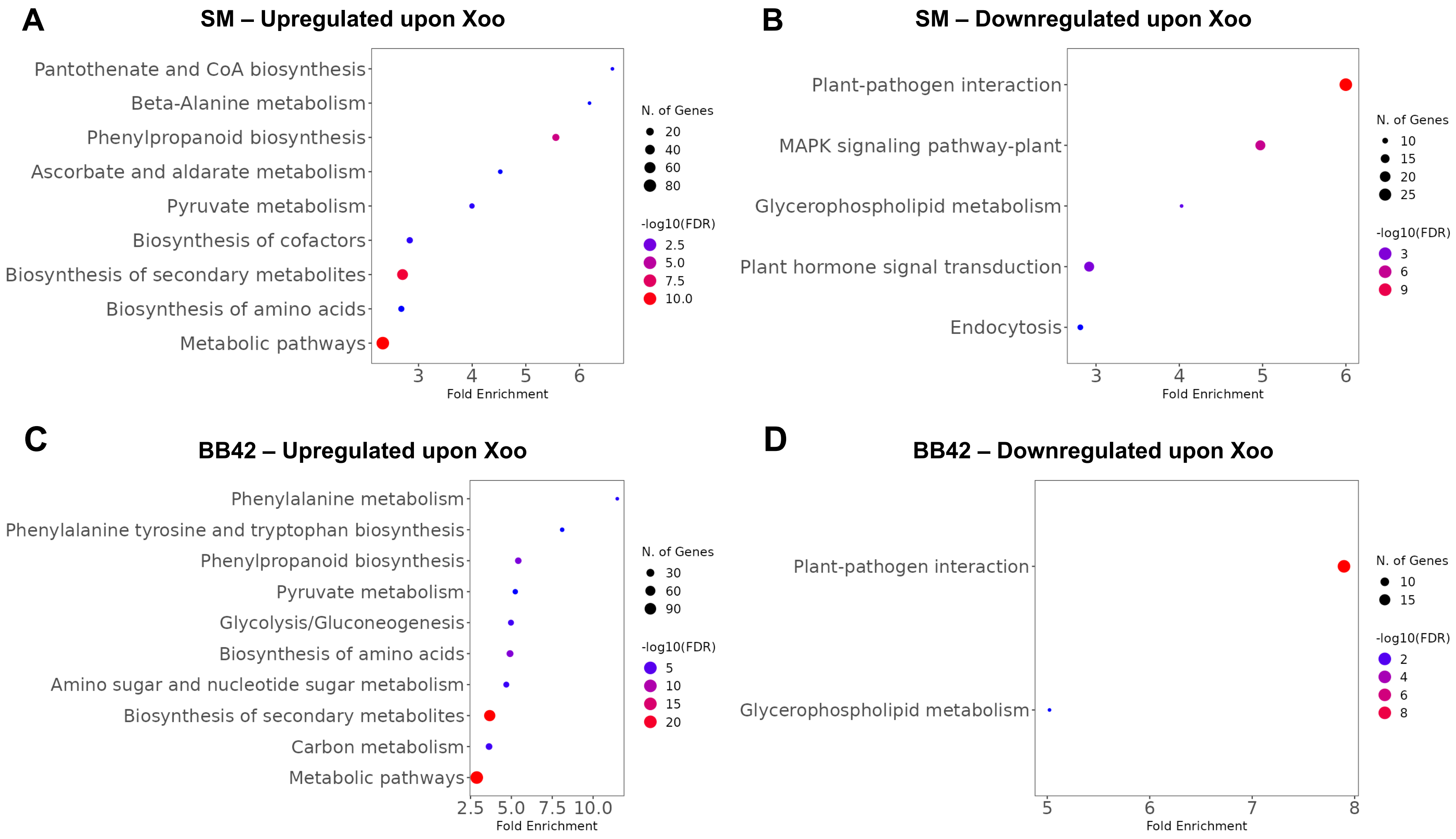


**Supplementary Figure S5:** Pathway analysis of the up and downregulated genes of Samba Mahsuri (SM) (A and B) and BB42 (C and D) upon *Xoo* treatment as compared to mock-treated samples.

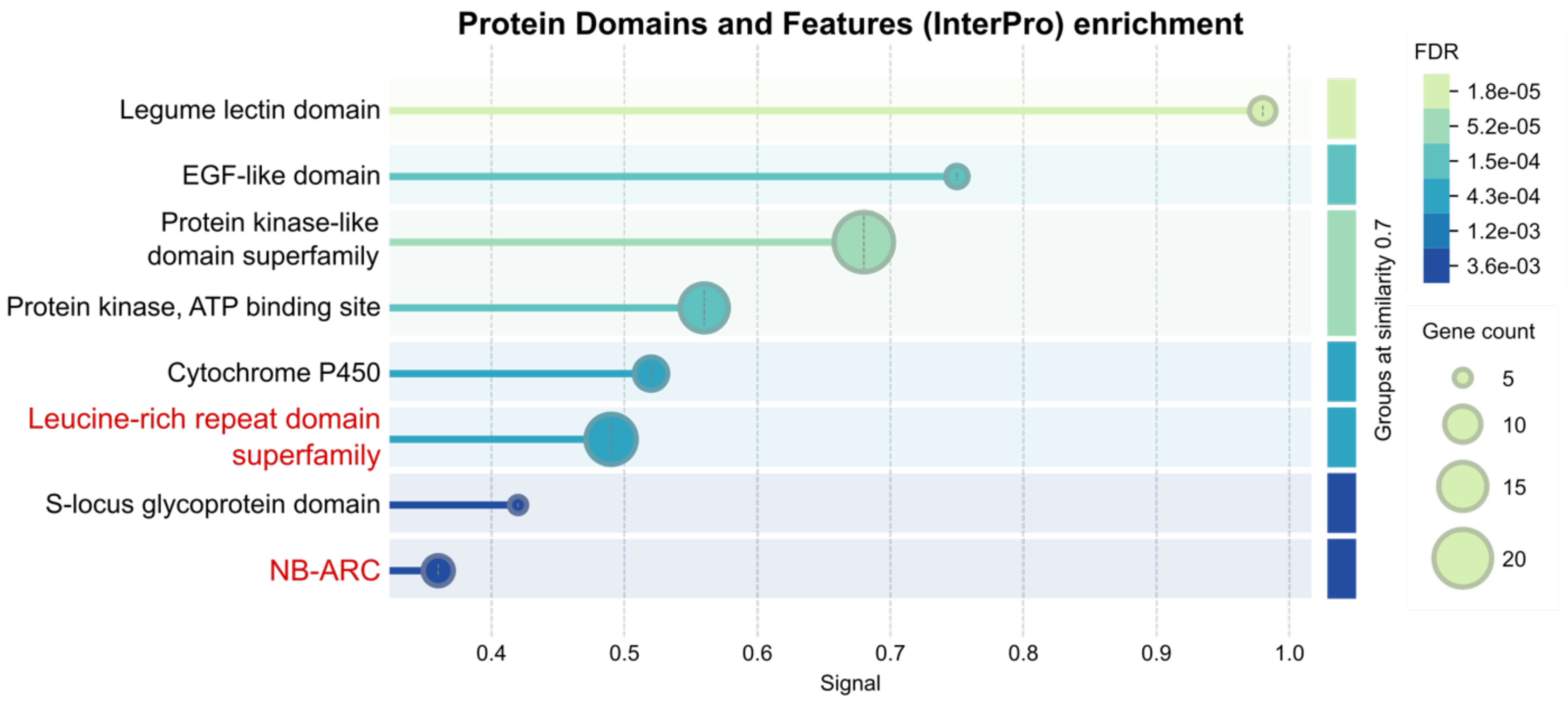


**Supplementary Figure S6:** Enrichment of protein domains among the upregulated genes in mock-treated BB42 as compared to mock-treated SM. Enrichment of LRR and NB-ARC features are highlighted in red. The enrichment analysis was performed using STRING v12.0.

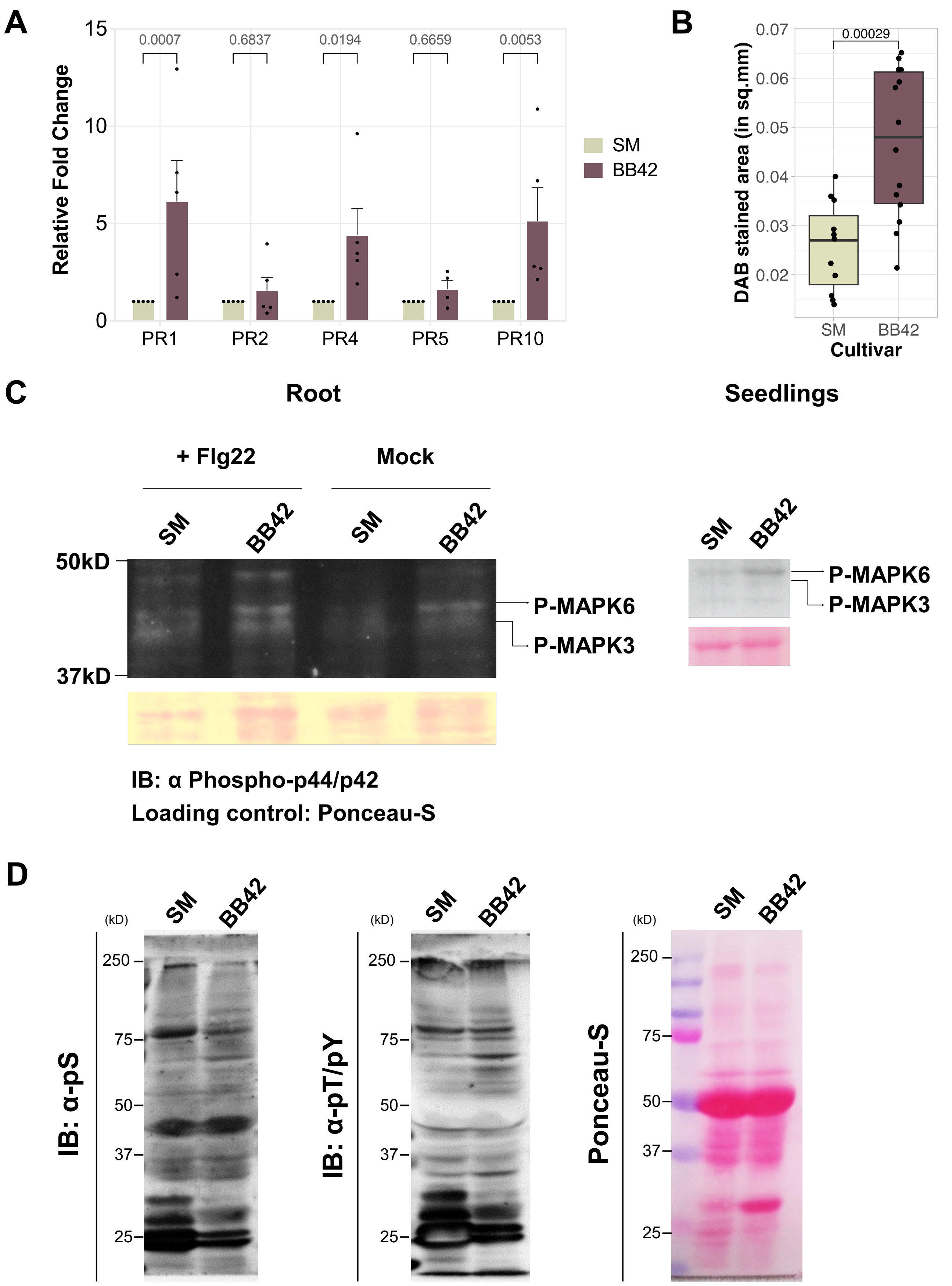

A
Relative Fold Change
0.0007
0.6837
0.0194
0.6659
0.0053
PR1
PR2
PR4
PR5
PR10
SM
BB42
B
DAB stained area (in sq.mm)
0.00029
Cultivar
C
Root
Seedlings
+ Flg22
Mock
50kD
37kD
P-MAPK6
P-MAPK3
IB: α Phospho-p44/p42
Loading control: Ponceau-S
D
(kD)
IB: α-pS
IB: α-pT/pY
Ponceau-S

**Supplementary Figure S7:** (A) Bar plots showing the relative difference in the expression levels of defense marker genes in Samba Mahsuri (SM) and BB42 seedlings. Fold change values were calculated using the $2^{-\Delta\Delta Ct}$ method. Bars indicate average fold change values in 5 independent experiments and the error bars represent standard error of mean. The numbers above the horizontal lines indicate *P*-values calculated using two-way ANOVA with Tukey's multiple comparisons test. $P<0.05$ is considered statistically significant. (B) Box plot showing the DAB-stained area measured using the ImageJ tool. Boxes represent the 25th to 75th percentile while the line within the box indicates the median. The whiskers extend to maximum and minimum data points that are within 1.5 times the interquartile range. The number above the horizontal line indicate *P*-values calculated using unpaired Student's *t*-test considering equal variance. $P<0.05$ is considered statistically significant. (C) Immunoblots of SM and BB42 roots (1-week old) and seedling (1-week old) samples probed using the anti-Phospho-erk1/2 antibody (1:2000 dilution) with or without Flg22 treatment and sampled after 15 minutes. Ponceau-S-stained blots are used as loading control blots. (D) Immunoblots of SM and BB42 leave samples probed using anti-Phosphoserine (ab9332; 1:1000; Abcam) and anti-Phosphothreonine/tyrosine (#9381; 1:3000; Cell Signaling Technologies) antibodies. Ponceau-S-stained blot is used as loading control blots. The immuno signals were visualized using HRP-conjugated anti-rabbit secondary antibody (ab98440; 1:20000; Abcam) and ECL Western substrate.

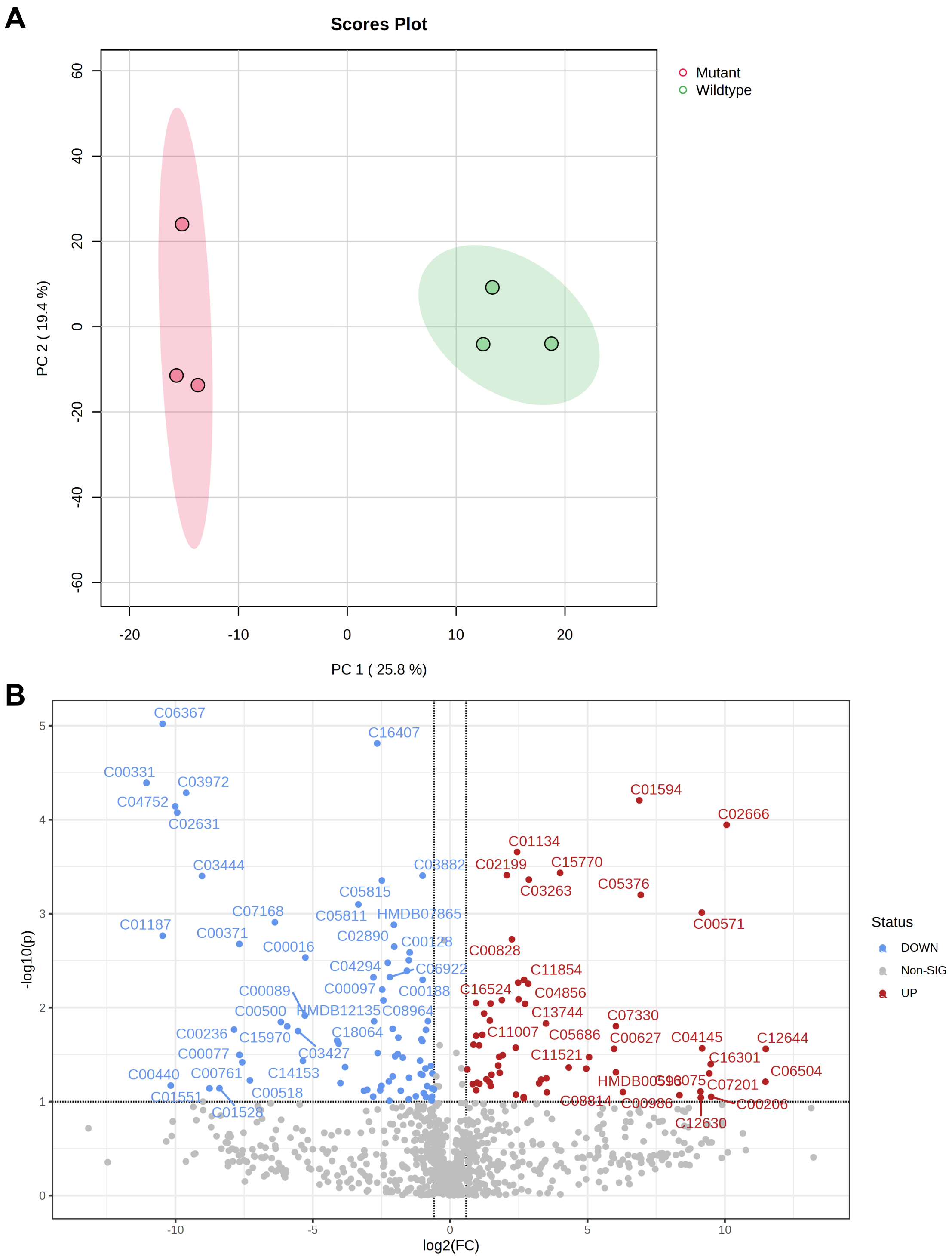


**Supplementary Figure S8:** (A) Principal Component Analysis of metabolite abundance in the matured leaves of SM and BB42. (B) Volcano plot showing the differentially accumulated metabolites in the leaves of SM and BB42.

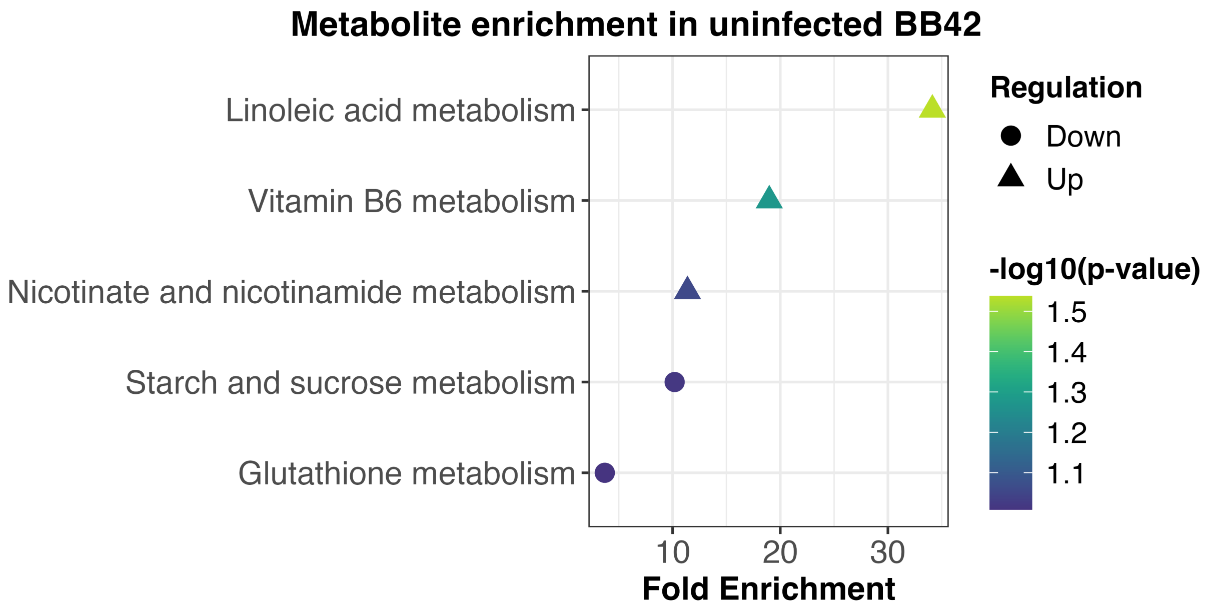


**Supplementary Figure S9:** KEGG pathway analysis of the differentially accumulated metabolites in the leaves of SM and BB42.

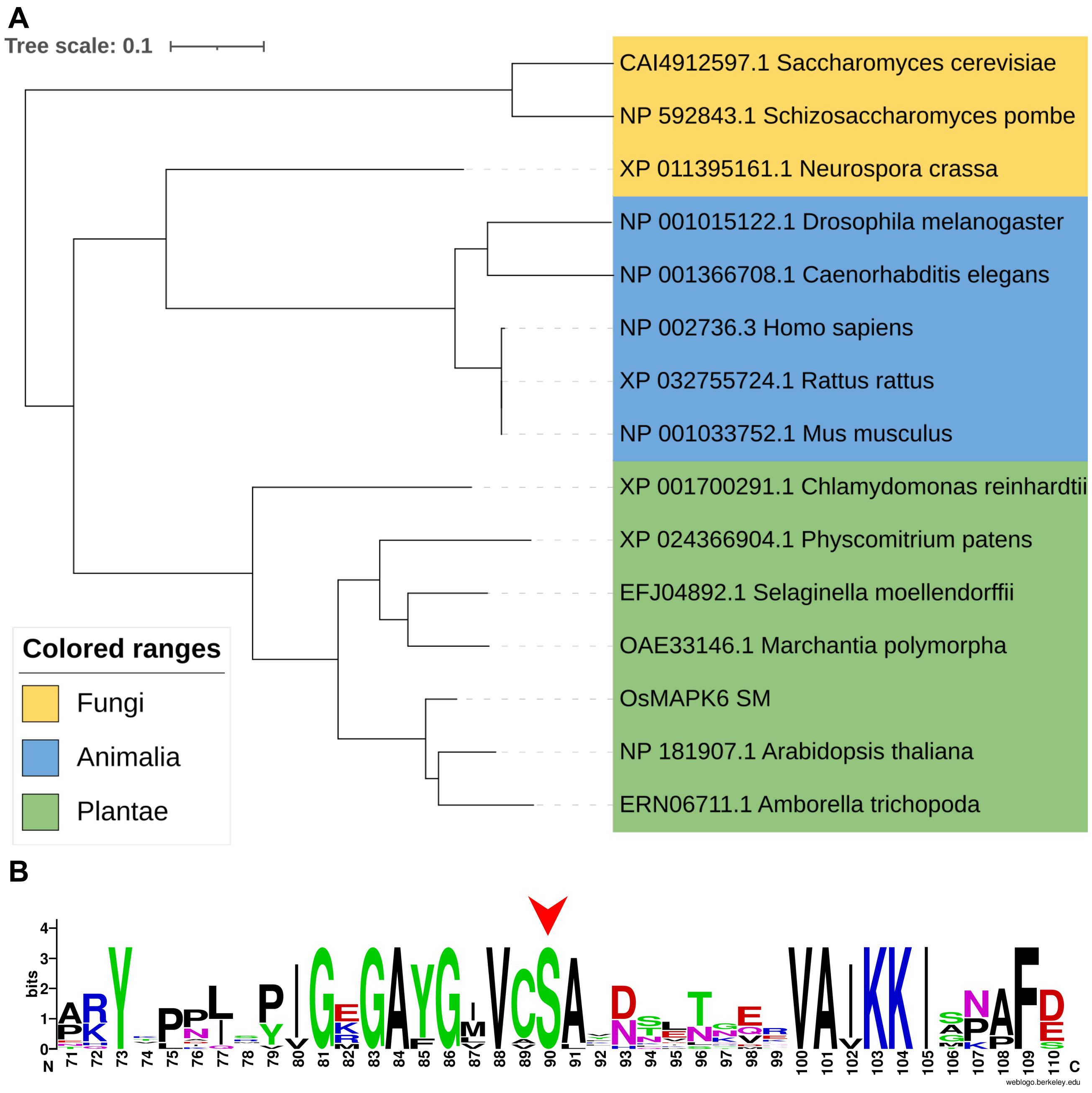


**Supplementary Figure S10:** (A) A maximum likelihood phylogenetic tree constructed using the MAPK6 homologues from representative species of the kingdoms Plantae, Animalia, and Fungi. The sequences were obtained using NCBI-BLASTp search with OsMAPK6-SM as query. The sequences were subjected to phylogenetic analysis with the "one click" mode of Phylogeny.fr programme and the tree file was annotated using iTOL (https://itol.embl.de). (B) Logo of the multiple sequence alignment of the sequences from (A), spanning the Serine residue of interest (red arrowhead), showing that the residue is invariant. The logo was generated using the WebLogo application (https://weblogo.berkeley.edu).

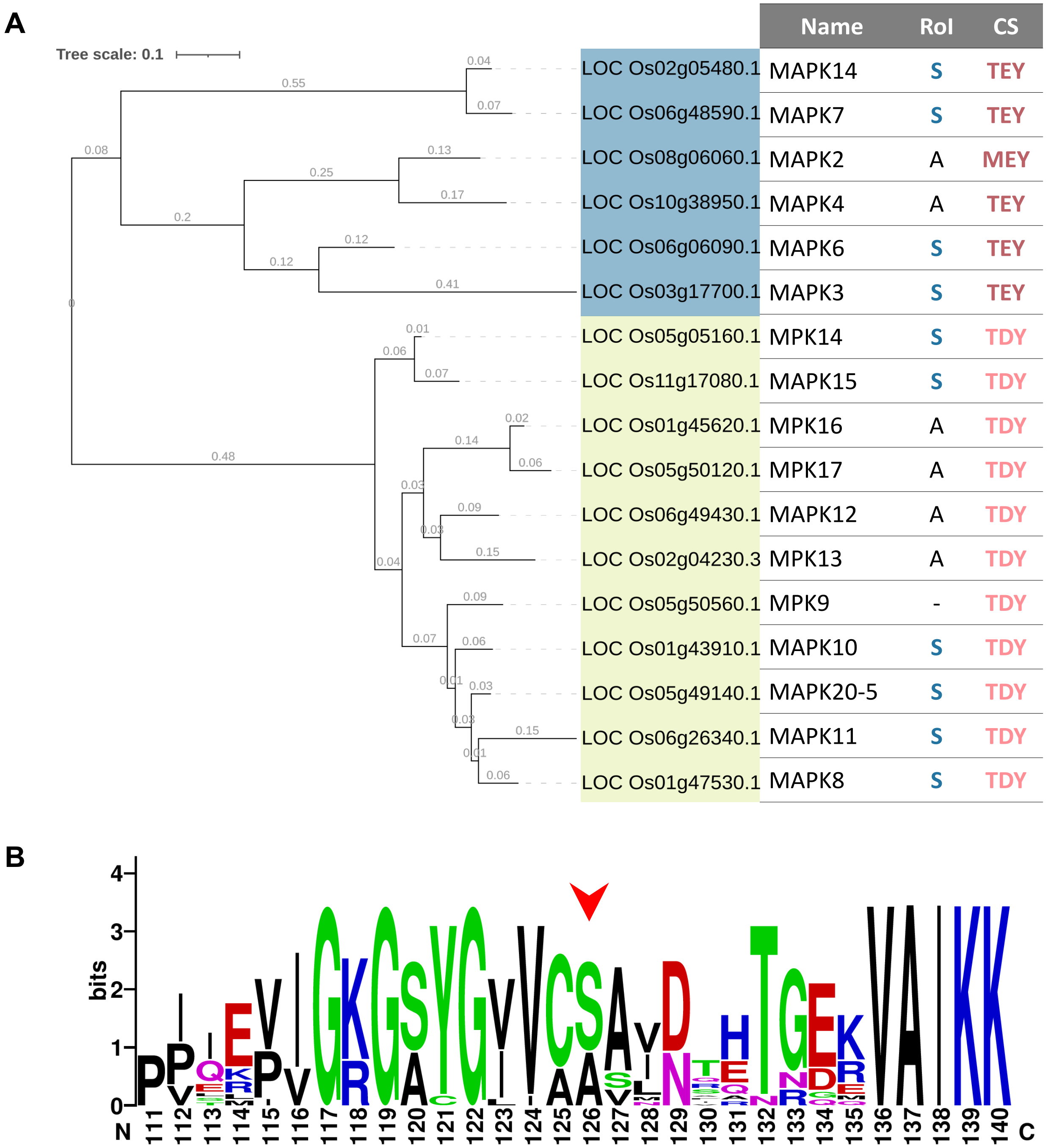


**Supplementary Figure S11:** (A) A maximum likelihood phylogenetic tree of the seventeen rice MAP Kinases. The sequence IDs were obtained from Yang *et al.*, (2015) and the sequences from RGAP database Hamilton *et al.*, (2025). The sequences were subjected to phylogenetic analysis with the "one click" mode of Phylogeny.fr programme and the tree file was annotated using iTOL (https://itol.embl.de). RoI - residue of interest; CS - Catalytic Site. "-" indicates the absence of the residue of interest. (B) Logo of the multiple sequence alignment of the sequences from (A), spanning the Serine residue of interest (red arrowhead). The logo was generated using the WebLogo application (https://weblogo.berkeley.edu).
