## Supplementary material for "A novel allele of *Mitogen Activated Protein Kinase 6* is linked to disease resistance and constitutive immunity in rice": Table S

### Supplementary Tables

#### Supplementary Table 1 to 8

A novel allele of *Mitogen Activated Protein Kinase 6* is linked to disease resistance and constitutive immunity in rice.

Gokulan C.G.[1,6,†], Mohammed Jamaloddin[1,†], Deepti Rao[1], Namami Gaur[1,2], Amita Rani Supriyo[1], Nisha Sao[1], Deepak Niranjan[1,7], Shrish Tiwari[1], Gouri S. Laha[3], Kalyani M. Barbadikar[3], Raman Meenakshi Sundaram[3], Sheshu Madhav Maganti[3,4,*], Hitendra K. Patel[1,5,*], Ramesh V. Sonti[1,7,*]

**Supplementary Table S1**: List of bacterial strains used in this study.

| Strains | Organism | Source |
|---|---|---|
| BXO1 | *Xanthomonas oryzae* pv. *oryzae* | Lab stock |
| BXO8 | *Xanthomonas oryzae* pv. *oryzae* | Lab stock |
| IXO631 | *Xanthomonas oryzae* pv. *oryzae* | Lab stock |
| IXO685 | *Xanthomonas oryzae* pv. *oryzae* | Lab stock |
| IXO1088 | *Xanthomonas oryzae* pv. *oryzae* | Lab stock |
| IXO1221 | *Xanthomonas oryzae* pv. *oryzae* | Lab stock |
| CHN/J | *Xanthomonas oryzae* pv. *oryzae* | IIRR |
| FZB-09-5 | *Xanthomonas oryzae* pv. *oryzae* | IIRR |
| LVD-09-1-1 | *Xanthomonas oryzae* pv. *oryzae* | IIRR |
| RPR-09-1 | *Xanthomonas oryzae* pv. *oryzae* | IIRR |
| BXOR1 | *Xanthomonas oryzae* pv. *oryzicola* | Lab stock |

**Supplementary Table S2**: List of primers used in this study.

| Primer name | Sequence (5' to 3') | Purpose/Gene ID |
|---|---|---|
| MAPK6-IFP | CGTCTCCGAGTTGAGCGCAGA | Tetra-ARMS-PCR |
| MAPK6-IRP | GATTTTTGGATTGGGATGGCATCC | |
| MAPK6-OFP | GACATTCTCGTGGTCCATGTGGC | |
| MAPK6-ORP | GAGGTCACCGCCAAGTACAAGCC | |
| SM-dCAPS-FP | GGAGGGAATTCCGTGGTCGAAAC | dCAPS |
| SM-dCAPS-RP | GGATTTTTGGATTGGGATGGGAGC | |
| BB42-dCAPS-FP | CCCGTCTCCGAGTTGAGCCCCG | |
| BB42-dCAPS-RP | TCGGGAACGTGTTCGAGGTCACC | |
| PR1-RT-FP | ACTCTGTACACTGTGATGGATGCG | qPCR/LOC_Os07g03730 |
| PR1-RT-RP | TACACGAGTCTTGCACGTTGGC | |
| PR2-RT-FP | ATTGTACAGAGGGCTTGGCTTGC | qPCR/LOC_Os05g31140 |
| PR2-RT-RP | CGCATCTGACAGACGTACACTTGG | |
| PR4-RT-FP | TGGCAAATGTCTCTCGGTGACG | qPCR/LOC_Os11g37960 |
| PR4-RT-RP | TGGCCATTCTGGTAGCCTTGAC | |
| PR5-RT-FP | ACGCCGCAAGTCATGTCCTAAAG | qPCR/LOC_Os12g43380 |
| PR5-RT-RP | AAACAATTGCACACGTGGTCGAG | |
| PR10-RT-FP | TGCGGGAGTCGGGAATACATAC | qPCR/LOC_Os12g36860 |
| PR10-RT-RP | CCAGCACCTCTGACTTTAGCAC | |
| GAPDH-RT-FP | AAGCCAGCATCCTATGATCAGATT | qPCR/LOC_Os04g40950 |
| GAPDH-RT-RP | CGTAACCCAGAATACCCTTGAGTTT | |
| Chr6_20706490_FP | TACGGGTGGTTTCACGTACGTGTC | To confirm the hybridity of the $F_1$ individuals. |
| Chr6_20706490_RP | CTACTCGAGGACGTGCCCCAAC | |
| Chr11_23983687_FP | CCCGTTGATCTCCCGAACAAATTGG | |
| Chr11_23983687_RP | GACCAGCTGCATGGTCCTTCAGAA | |

**Supplementary Table S3:** Statistics of the RNA-sequencing data generated in the study. B0 - BB42 0hpi/Mock; B24 - BB42 24hpi/*Xoo*; S0 - SM 0hpi/Mock; S24 - SM 24hpi/*Xoo*. The biological replicates of each treatment is represented as -1,-2,-3.

| Sample | Number of raw read pairs | Number of mapped reads | Number of uniquely mapped reads | Number of reads assigned to genes |
|---|---|---|---|---|
| B0-1 | 38484480 | 29709527 | 11845019 | 9816920 |
| B0-2 | 37092229 | 29681183 | 11597409 | 9625171 |
| B0-3 | 47424175 | 36709460 | 15508634 | 12793023 |
| B24-1 | 45482028 | 36234011 | 15646014 | 13044262 |
| B24-2 | 35781802 | 31527376 | 12526770 | 10441640 |
| B24-3 | 39309247 | 30918471 | 13534466 | 11447493 |
| S0-1 | 35498002 | 27691009 | 10319295 | 8315979 |
| S0-2 | 44941445 | 35406160 | 12403620 | 9859639 |
| S0-3 | 41068774 | 31758283 | 11327857 | 8971994 |
| S24-1 | 41495156 | 31978805 | 12536891 | 10238129 |
| S24-2 | 30432059 | 23599375 | 8939725 | 7303608 |
| S24-3 | 39895814 | 31303600 | 11483689 | 9123944 |

hpi: Hours post infection, *Xoo*: *Xanthomonas oryzae* pv. *oryzae*

**Supplementary Table S4:** Phenotype segregation analysis of bacterial blight resistance and susceptibility in two $F_2$ populations.

| **Population** | **Cross** | **Total** | **Susceptible (Median lesion length)** | **Resistant (Median lesion length)** | **Chi squared** | ***P*-value** | **R-bulk (n)** | **S-bulk (n)** |
|---|---|---|---|---|---|---|---|---|
| Population-1 | SM x BB42 | 244 | 191 (16.5 cm) | 53 (11.3 cm) | 2.14 | 0.1435 | 24 | 20 |
| Population-2 | BB42 x SM | 244 | 191 (15.6 cm) | 53 (9.5 cm) | 2.14 | 0.1435 | 29 | 20 |

SM: Samba Mahsuri, cm: Centimetre, R-bulk, Resistant bulk, S: Susceptible bulk, (n): Number of lines

**Supplementary Table S5:** Differential gene expression statistics upon various comparisons of the transcriptome data.

| S. No. | Source | Comparison | Upregulated | Downregulated | Total DEGs |
|---|---|---|---|---|---|
| 1. | SM | *Xoo* – 24hpi vs 0hpi | 778 | 1196 | **1974** |
| 2. | BB42 | *Xoo* – 24hpi vs 0hpi | 845 | 632 | **1477** |
| 3. | 0hpi (Mock) | BB42 vs SM | 193 | 617 | **810** |
| 4. | 24hpi (*Xoo*) | BB42 vs SM | 294 | 199 | **493** |

SM: Samba Mahsuri, hpi: Hours post infection, *Xoo*: *Xanthomonas oryzae* pv. *Oryzae*, DEGs: Differentially expressed genes

**Supplementary Table S6:** List of defense related genes and their fold change values in BB42 as compared to SM. NB-ARC/NLR encoding genes are in bold.

| Gene ID | Gene name | Description | log2FC |
|---|---|---|---|
| Os01g0116800 | | Similar to Receptor-like kinase. | 1.15 |
| Os01g0178900 | | **Nucleotide-binding LRR receptor (NLR) family protein** | 1.27 |
| Os02g0115700 | *OsCAT1A, OsCAT1, OsCATC* | Catalase A, Environmental stress response, Drought stress tolerance | 1.46 |
| Os02g0125800 | *DLN40, OsDLN40* | Transcription factor Tfb4 family protein. | 1.08 |
| Os02g0153700 | *OsLRK4, OsPSKR6* | **Leucine-rich repeat receptor-like kinase** | 1.26 |
| Os02g0227700 | *OsRLK5* | **LRR-receptor-like kinase (LRR-RLK) family protein** | 1.81 |
| Os02g0228000 | *OsRLCK70* | Protein kinase, catalytic domain containing protein. | 3.06 |
| Os02g0282000 | | **Nucleotide-binding LRR receptor (NLR) family protein, Resistance to bacterial blight** | 2.44 |
| Os02g0472566 | | Allergen V5/Tpx-1-related domain containing protein. | 1.18 |
| Os02g0504200 | | **Nucleotide-binding LRR receptor (NLR) family protein** | 1.46 |
| Os03g0225900 | *OsAOS2* | Allene oxide synthase (CYP74A2), Biosynthesis of jasmonic acid (JA) | 4.09 |
| Os04g0193950 | *OsLR10* | **Disease resistance protein domain containing protein.** | 1.33 |
| Os04g0227200 | | **LRR-receptor-like kinase (LRR-RLK) family protein** | 1.78 |
| Os04g0288500 | | Protein kinase, catalytic domain containing protein. | 1.26 |
| Os04g0370100 | *OsWAK46, OsWAK2L* | Similar to OSIGBa0107E14.7 protein. | 2.92 |
| Os06g0328400 | | Protein kinase, catalytic domain containing protein. | 1.16 |
| Os06g0602500 | *OsNRFG6, OsSDRLK38* | S-Domain receptor like kinase-38, Response to drought, chilling and Xanthomonas oryzae pv. oryzae in tolerant rice genotypes, Response to submergence | 1.29 |
| Os06g0614100 | *OsbZIP49, OsbZIP50* | A member of the TGA family of basic leucine zipper transcription factors, Control of tiller angle and plant architecture, Regulation of auxin homeostasis | 1.85 |
| Os07g0117300 | | **NB-ARC domain containing protein.** | 1.97 |
| Os07g0130800 | | Protein kinase, catalytic domain containing protein. | 2.47 |

| | | | |
|---|---|---|---|
| Os07g0131375 | | Protein kinase, catalytic domain containing protein. | 1.07 |
| Os07g0283050 | | Concanavalin A-like lectin/glucanase, subgroup domain containing protein. | 3.76 |
| Os08g0174800 | | Similar to Benzothiadiazole-induced somatic embryogenesis receptor kinase 1. | 2.46 |
| Os08g0360300 | *OsSARD1* | Calmodulin binding protein-like domain containing protein. | 1.60 |
| Os09g0268000 | | Similar to lectin-like receptor kinase 7. | 1.51 |
| Os09g0268100 | | Similar to lectin-like receptor kinase 7. | 5.33 |
| Os09g0561600 | *OsWAK91, OsDEES1* | Wall-associated receptor-like kinase, Regulation of early embryo sac development, Female gametophyte development, Positive regulation of rice blast resistance | 1.19 |
| Os10g0142600 | *OsWAK101, OsRLCK292* | Protein kinase, catalytic domain containing protein. | 1.03 |
| Os10g0180800 | *OsWAK112, OsWAK112d* | Wall-associated kinase, Negative regulation of rice blast resistance | 1.16 |
| Os11g0133100 | *OsSDRLK58* | S-Domain receptor like kinase-58, Response to Rice Stripe Virus | 1.02 |
| Os11g0550100 | | **Similar to NB-ARC domain containing protein, expressed.** | 1.09 |
| Os11g0550500 | | **Similar to LZ-NBS-LRR class RGA.** | 1.01 |
| Os11g0673600 | *YR2, YR6, YR9, YR10, YR21, YR23* | **Similar to NB-ARC domain containing protein, expressed.** | 1.09 |
| Os11g0674400 | | **Similar to NB-ARC domain containing protein, expressed.** | 1.02 |
| Os11g0676100 | | **Similar to NB-ARC domain containing protein, expressed.** | 3.36 |
| Os11g0676500 | | Hypothetical conserved gene. | 1.11 |
| Os11g0677000 | | **Similar to NB-ARC domain containing protein, expressed**. | 1.78 |
| Os12g0130300 | *OsSDRLK57* | S-Domain receptor like kinase-57, Response to drought in tolerant genotype Nagina 22, Response to Rice Stripe Virus | 3.23 |
| Os12g0555300 | *OsPR10C* | Pathogenesis-related protein PR-10c, Nonfunctional pseudogene | 1.70 |
| Os12g0556500 | | Calmodulin binding protein-like family protein. | 1.06 |
| Os12g0611100 | *OsRLCK373* | Similar to serine/threonine-protein kinase receptor. | 3.68 |

SM: Samba Mahsuri, log2FC: log2 fold change

**Supplementary Table S7:** List of JASMONATE ZIM-DOMAIN (JAZ) protein-coding genes and their fold change values in BB42 in comparison to SM.

| Gene ID | Gene name | Description | log2FC |
|---|---|---|---|
| Os03g0402800 | *OsTIFY10A* | TIFY family protein, JASMONATE-ZIM domain (JAZ) protein, JA signalling, Regulation of spikelet development, Regulation of coleoptile length under submergence | -2.81 |
| Os07g0615200 | *OsTIFY10B* | JASMONATE ZIMDOMAIN (JAZ) protein, Regulation of coleoptile length under submergence | -2.24 |
| Os09g0439200 | *OsTIFY10c* | Jasmonate ZIM-domain protein, Jasmonate-induced resistance to bacterial blight, Repressor of jasmonic acid signalling | -2.14 |
| Os03g0180800 | *OsTIFY11a* | TIFY domain-containing transcriptional regulator, Salt and dehydration stress tolerance | -3.62 |
| Os03g0181100 | *OsTIFY11b* | TIFY family protein, Jasmonate ZIM-domain (JAZ) protein, miR156-target gene, Signal transduction induced by hydrogen peroxide | -4.41 |
| Os03g0180900 | *OsTIFY11c* | Jasmonate ZIM-domain containing protein, Transcriptional repressor of JA signalling, Regulation of phosphate starvation responses, Pi homeostasis | -2.50 |
| Os10g0392400 | *OsTIFY11d* | Tify domain containing protein. | -3.62 |
| Os10g0391400 | *OsTIFY11e* | Jasmonate ZIM-domain (JAZ) protein, TIFY family protein, Negative regulation of JA signal transduction pathway, Activation of hypersensitive cell death | -6.02 |

SM: Samba Mahsuri, log2FC: log2 fold change

**Supplementary Table S8:** List of genes and their differential expression related to various components of brassinosteroid (BR) pathway.

| Gene ID | Gene name | Log2FC | Role |
|---|---|---|---|
| Os01g0705700 | *bHLH10/OsICE1* | -3.75 | Response to BR stimulus |
| Os08g0490000 | *bHLH31/OsBIM2* | -2.37 | BR signaling |
| Os01g0625900 | *OFP2* | -2.09 | Response to BR stimulus |
| Os10g0575000 | *bHLH9/MYC2* | -2.05 | BR signaling |
| Os08g0465000 | *HOX27* | -2.03 | Response to BR stimulus |
| Os01g0757200 | *GA2OX3* | -1.99 | BR signaling |
| Os05g0343400 | *WRKY53* | -1.52 | Response to BR stimulus |
| Os07g0569100 | *REM4.1* | -1.35 | BR signaling |
| Os07g0597200 | *BIN1* | -1.35 | BR signaling |
| Os03g0227700 | *DWARF4* | -1.24 | BR biosynthesis |
| Os10g0558700 | *ODD11* | 1.65 | Response to BR stimulus |
| Os07g0650600 | *BLE2* | 1.69 | Response to BR stimulus |

log2FC: log2 fold change
